# The SUMO-interacting Motif in NS2 promotes adaptation of avian influenza virus to mammals

**DOI:** 10.1101/2022.12.11.519849

**Authors:** Liuke Sun, Huihui Kong, Mengmeng Yu, Zhenyu Zhang, Haili Zhang, Lei Na, Yuxing Qu, Yuan Zhang, Hualan Chen, Xiaojun Wang

**Author notes:** Co-first author (Liuke Sun, Huihui Kong, Mengmeng Yu).

## Abstract

Species differences in the host factor ANP32A/B result in the restriction of avian influenza virus polymerase (vPol) in mammalian cells. Efficient replication of avian influenza viruses in mammalian cells often requires adaptive mutations, such as PB2-E627K, to enable the virus to utilize mammalian ANP32A/B. However, the molecular basis for the productive replication of avian influenza viruses without prior adaptation in mammals remain poorly understood. We show that influenza A virus protein NS2 help to overcome mammalian ANP32A/B-mediated restriction to avian vPol activity by promoting avian vRNP assembly and enhancing mammalian ANP32A/B-vRNP interactions. A conserved SUMO interaction motif (SIM) in NS2 is required for its avian polymerase-enhancing properties. We also demonstrate that disrupting SIM integrity in NS2 impairs avian influenza virus replication and pathogenicity in mammalian hosts, but not in avian hosts. Our results identify NS2 as a cofactor in the adaptation process of avian influenza viruses to mammals.

## INTRODUCTION

Host restriction limits cross-species transmission of avian influenza virus (AIVs) from migratory aquatic birds to mammals(Long et al., 2019b). However, some highly pathogenic avian influenza strains (e.g., H5N1 and H7N9) have overcome this host barrier and are able to infect humans, resulting in a serious disease with a high mortality rate, and posing a huge threat to human welfare(Li and Chen, 2021; Li et al., 2014a; Neumann et al., 2010; Su et al., 2017; Zhang et al., 2013a; Zhou et al., 2013). In addition, it is not uncommon for a human host to be infected with multiple different subtypes of avian influenza viruses, for example, H9N2, H6N1, H7N7, H10N8, H7N3 and H5N6, concurrently(Belser et al., 2013; Chen et al., 2014; Gu et al., 2022; Jonges et al., 2014; Pan et al., 2018; Wei et al., 2013). The avian influenza virus must overcome multiple host range barriers to effectively infect and spread among mammals. A major species barrier is the restriction of avian vPol in mammalian cells(Long et al., 2016; Mehle and Doudna, 2008), which is the driving force for the emergence of adaptive mutations(Liang et al., 2019). Two most well-known mammalian adaptation mutation, E627K and D701N in PB2, increases the replication, pathogenicity, and transmittance of avian influenza viruses in mammals(Almond, 1977; Gabriel et al., 2005; Gao et al., 2009; Hatta et al., 2001; Mok et al., 2014; Steel et al., 2009; Subbarao et al., 1993; Van Hoeven et al., 2009). Recent studies have identified that species-specific differences in proteins of the acidic nuclear phosphoprotein 32 (ANP32) family determine the restriction of avian vPol in mammalian cells(Long *et al*., 2016).

The host factor ANP32A/B regulates the synthesis of vRNA from influenza A virus (IAVs) cRNA(Sugiyama et al., 2015). Compared with human ANP32A/B (huANP32A/B), chicken ANP32A (chANP32A) has a unique insertion of 33 amino acids that enables it to efficiently support the vPol activity of both avian and human-adapted influenza viruses. However, huANP32A/B only supports the vPol activity of human-adapted influenza virus, and cannot support that of the avian influenza virus, due to the absence of this insertion(Long *et al*., 2016). In addition, further studies have revealed that the avian hosts carry up three chANP32A isoforms. These three isoforms differ only in the composition of the 33 amino acid inserts, but support the activity of avian-signature vPol to differing extents(Baker et al., 2018; Domingues et al., 2019). A hydrophobic SUMO interaction motif (SIM)-like sequence in the 33-amino acid insertion of chANP32A is required to selectively support avian vPol activity(Domingues and Hale, 2017), suggesting that SUMO-dependent function is important for the function of avian vPol. In addition, results from our lab and others have demonstrated that huANP32A and huANP32B equally support the polymerase activity of human-adapted influenza A virus, and that loss of huANP32A and huANP32B simultaneously results in failure of influenza A virus replication(Staller et al., 2019; Zhang et al., 2019). Chicken ANP32B (chANP32B) has natural mutations at positions 129 and 130, and it has lost its ability to support the activity of influenza A virus polymerase(Long et al., 2019a; Zhang *et al*., 2019). Swine ANP32A (swANP32A) is able to support avian vPol activity to a greater extent than other mammalian ANP32A/B, as a result of a recently discovered unique 106V/156S signature, which explains why pigs can act as intermediate hosts for the cross-species transmission of avian influenza virus(Peacock et al., 2020; Zhang et al., 2020a). In addition, human ANP32 proteins, including ANP32A, ANP32B and ANP32E, have recently been identified as cofactors of influenza B virus polymerase(Zhang et al., 2020b).

Mammalian ANP32A/B proteins, with the exception of swANP32A, provide poor support for avian vPol activity, so to achieve high levels of replication in mammalian cells, avian influenza viruses require mutations to adapt to mammalian ANP32A/B(Bi et al., 2019; Long *et al*., 2016; Peacock *et al*., 2020; Zhang *et al*., 2020a; Zhang *et al*., 2019). However, some human isolates of avian influenza viruses H9N2, H5N1 and H7N9 were able to establish productive infections in humans without the emergence of previously identified adaptive mutations(Le et al., 2010; Manz et al., 2016; Song and Qin, 2020). Avian isolates of multiple subtypes of avian influenza virus are even able to establish productive infections in mammalian hosts and are even pathogenic for mice without prior adaptation(Cui et al., 2022; Guan et al., 2019; Guo et al., 2021; Li et al., 2014b; Li et al., 2010; Liang *et al*., 2019; Shi et al., 2017; Shi et al., 2018; Yin et al., 2021; Zhou *et al*., 2013). These studies suggest that the polymerases of certain subtypes of avian influenza viruses can use mammalian ANP32A/B to some extent for productive replication in the process of cross-species transmission into mammals. This contradicts the conclusions drawn from cell-based vPol reconstitution assays that mammalian ANP32A/B poorly support avian vPol activity(Long *et al*., 2016; Peacock *et al*., 2020; Zhang *et al*., 2020a; Zhang *et al*., 2019). Indeed, it has been demonstrated that during transcription and replication, the regulatory events that occur in the IAV replication process are quite different from those that occur in cell-based vPol reconstitution assays(Robb et al., 2009). Most importantly, with the exceptions of the trimeric polymerase complex and NP, the contributions of other IAV proteins produced during viral replication to avian vPol activity are largely unknown. In this study, we show that the influenza A virus NS2 promotes avian-signature vPol activity supported by mammalian ANP32A/B, but not chANP32A. NS2 has a relatively limited effect on the activity of mammalian-signature vPol supported by both mammalian ANP32A/B and chANP32A. NS2 proteins derived from influenza A viruses isolated from a range of hosts are conserved in this avian-signature polymerase-enhancing function. In addition, we demonstrate that NS2 exerts this function by facilitating the assembly of avian vRNP and enhancing the interaction of mammalian ANP32A/B with avian vRNP in mammalian cells. Most importantly, NS2 harbors a highly conserved SUMO-interacting motif (SIM), which is required for its avian-signature polymerase-enhancing properties. Disruption of the integrity of this SIM in NS2 has deleterious effects on the replication and pathogenicity of avian influenza viruses in mammalian hosts, but not in avian hosts. Taken together, our data suggest a model in which the adaptation of avian-signature vPol to mammalian ANP32A/B relies on the help of the SIM supplied by IAV-encoded NS2 protein, providing novel insights into the adaptation of avian influenza viruses to mammals.

## RESULTS

### Regulation of the activity of the avian H9N2 virus vPol by NS2

Host factor ANP32 proteins determine the function of IAVs polymerase, and mammalian cells are non-permissive for AIVs vPol because of restriction from their ANP32A/B(Baker *et al*., 2018; Bi *et al*., 2019; Domingues *et al*., 2019; Domingues and Hale, 2017; Long *et al*., 2016; Long *et al*., 2019a; Peacock *et al*., 2020; Staller *et al*., 2019; Zhang *et al*., 2020a; Zhang *et al*., 2019). However, many H9N2, H5N1, or H7N9 AIVs can establish productive infection in humans without prior adaptation(Guo *et al*., 2021; Le *et al*., 2010; Manz *et al*., 2016; Song and Qin, 2020; Yin *et al*., 2021). To investigate the mechanism by which the avian vPol can function efficiently in mammal cells, we used *huANP32A&B&E* triple-knockout HEK293T cells (TKO)(Zhang *et al*., 2020b) to establish an ANP32-dependent mini-replicon vPol reporter system. We found that huANP32A/B and canine ANP32A/B (caANP32A/B) poorly supported the vPol activity of the avian influenza viruses A/chicken/Zhejiang/B2013/2012(H9N2_ZJ12_) (Figure 1A), which was consistent with previous results(Long *et al*., 2016; Peacock *et al*., 2020; Zhang *et al*., 2020a; Zhang *et al*., 2019). However, when the replication ability of H9N2_ZJ12_ in mammalian cells was evaluated, the results suggested that the H9N2_ZJ12_ virus could establish productive replication in mammalian cell lines, including A549 cells, HEK293T cells and MDCK cells (Figures 1B-D). These data indicate that huANP32A/B or caANP32A/B support the vPol activity of the avian H9N2_ZJ12_ virus to a degree sufficient to support productive replication under infection conditions, which is inconsistent with the results of vPol reconstitution assays in TKO cells (Figure 1A).

**Figure 1.**
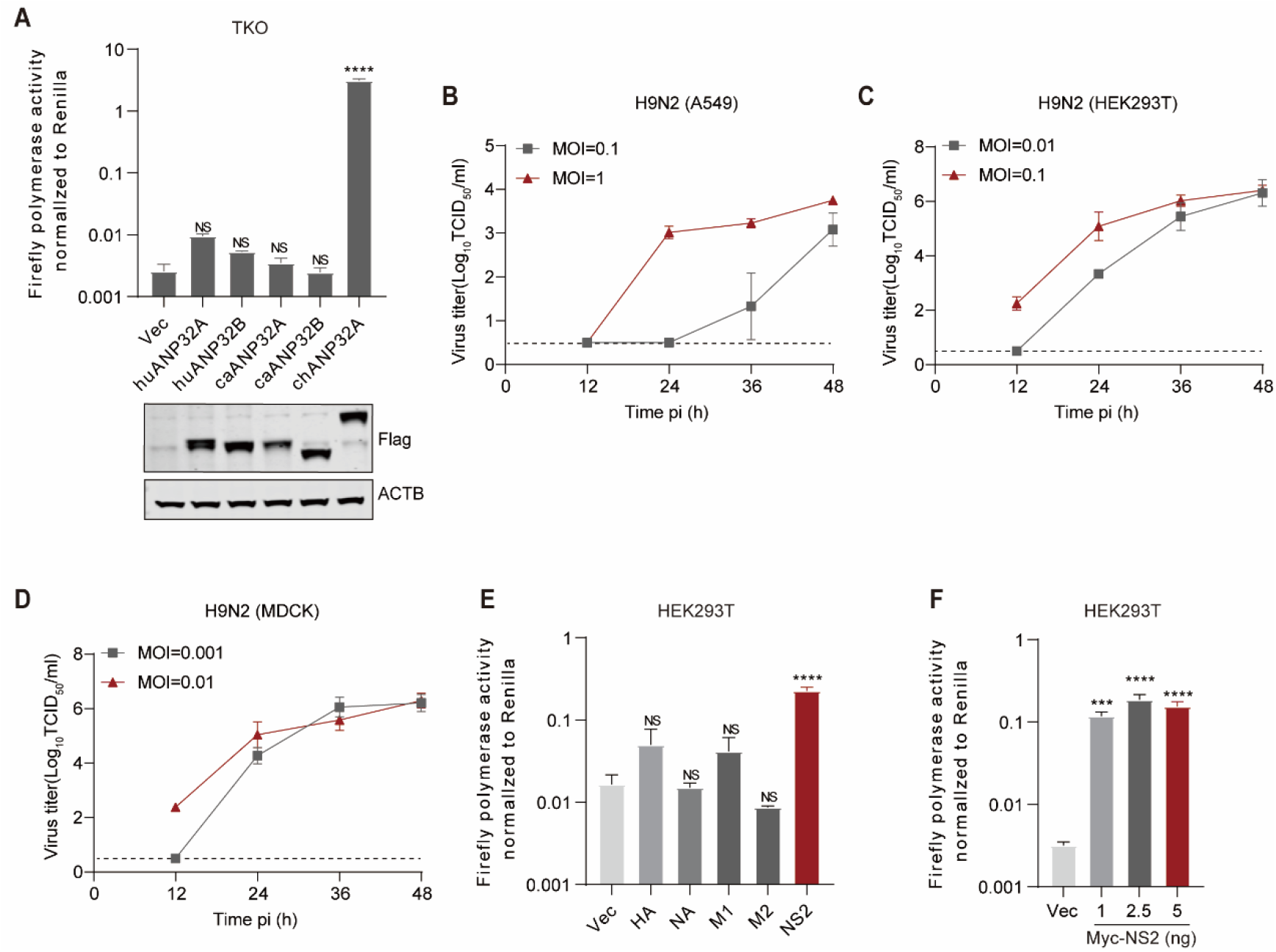
NS2 enhances H9N2 vPol activity in human cells. **A:** vPol reconstitution assay in TKO cells to compare the effect of each ANP32-Flag construct on avian H9N2 vPol activity. The accompanying western blots show expression of Flag-tagged ANP32 constructs. **B-D:** Viral growth kinetics of avian H9N2_ZJ12_ virus in the indicated cell lines. A549 cells (B), HEK293T cells (C) and MDCK cells (D) were infected with H9N2_ZJ12_ virus, and virus titers were determined in MDCK cells stably expressing chANP32A using TCID_50_ assays. **E:** vPol reconstitution assay to compare the effect of different influenza viral proteins on H9N2 vPol activity in HEK293T cells. **F:** vPol reconstitution assay in HEK293T cells to compare the effect of increasing doses of H9N2-NS2 on H9N2 vPol activity. In (A) -(F), bars represent mean values of the replicates within one representative experiment (n = 3, ± SD). Vec, empty vector control. Significance was determined with one-way ANOVA followed by a Dunnett’s multiple comparisons test (\*\*\**p* < 0.001; \*\*\*\**P* < 0.0001; NS, not significant).

As the vPol reconstitution assay involves only the ribonucleoproteins (RNP) required for viral transcription and replication, we hypothesized that besides these RNP proteins, other viral proteins produced under infection conditions could regulate the supporting function of mammalian ANP32A/B towards avian vPol activity and could account for the efficient replication of avian influenza virus in mammalian cells. To investigate this hypothesis, a vPol reconstitution assay was performed in HEK293T cells to assess whether other viral proteins produced during infection process, including HA, NA, M1, M2 and NS2, have an impact on the vPol activity of the H9N2_ZJ12_ virus. As shown in Figure 1E, H9N2-NS2 significantly enhanced the vPol activity of avian H9N2_ZJ12_ virus in HEK293T cells. In addition, we found that this enhancing effect of NS2 towards the vPol activity of the avian H9N2_ZJ12_ virus was dose-dependent (Figure 1F). Together, these data identified NS2 as a positive regulator of the vPol activity of the avian H9N2_ZJ12_ virus in human cells.

### Influenza A virus NS2 selectively promotes avian-signature vPol activity when supported by huANP32A/B, but not by chANP32A

We and other labs independently found that both huAP32A and huANP32B play important roles in supporting the vPol activity, and both exhibited a limited supporting function towards avian vPol activity(Peacock *et al*., 2020; Staller *et al*., 2019; Zhang *et al*., 2020a; Zhang *et al*., 2019). We next evaluated the effect of H9N2-NS2 on the avian and mammalian-signature (PB2^627K^) vPol activity of the H9N2_ZJ12_ virus supported by ANP32 proteins. To do this, we established an ANP32-dependent vPol reconstitution assay in TKO cells. We found that NS2 dramatically enhanced the avian H9N2 vPol activity when supported by huANP32A (∼47.01-fold increase) or huANP32B (∼105.99-fold increase), but not by chANP32A. However, NS2 had limited effects on the activity of the mammalian-signature H9N2 (PB2^627K^) vPol when supported by huANP32A, huANP32B or chANP32A (Figures 2A-C). We obtained consistent results using the NS2 and vPol from the genetic background of the avian influenza H7N9_AH13_ virus (Figures 2D-F).

**Figure 2.**
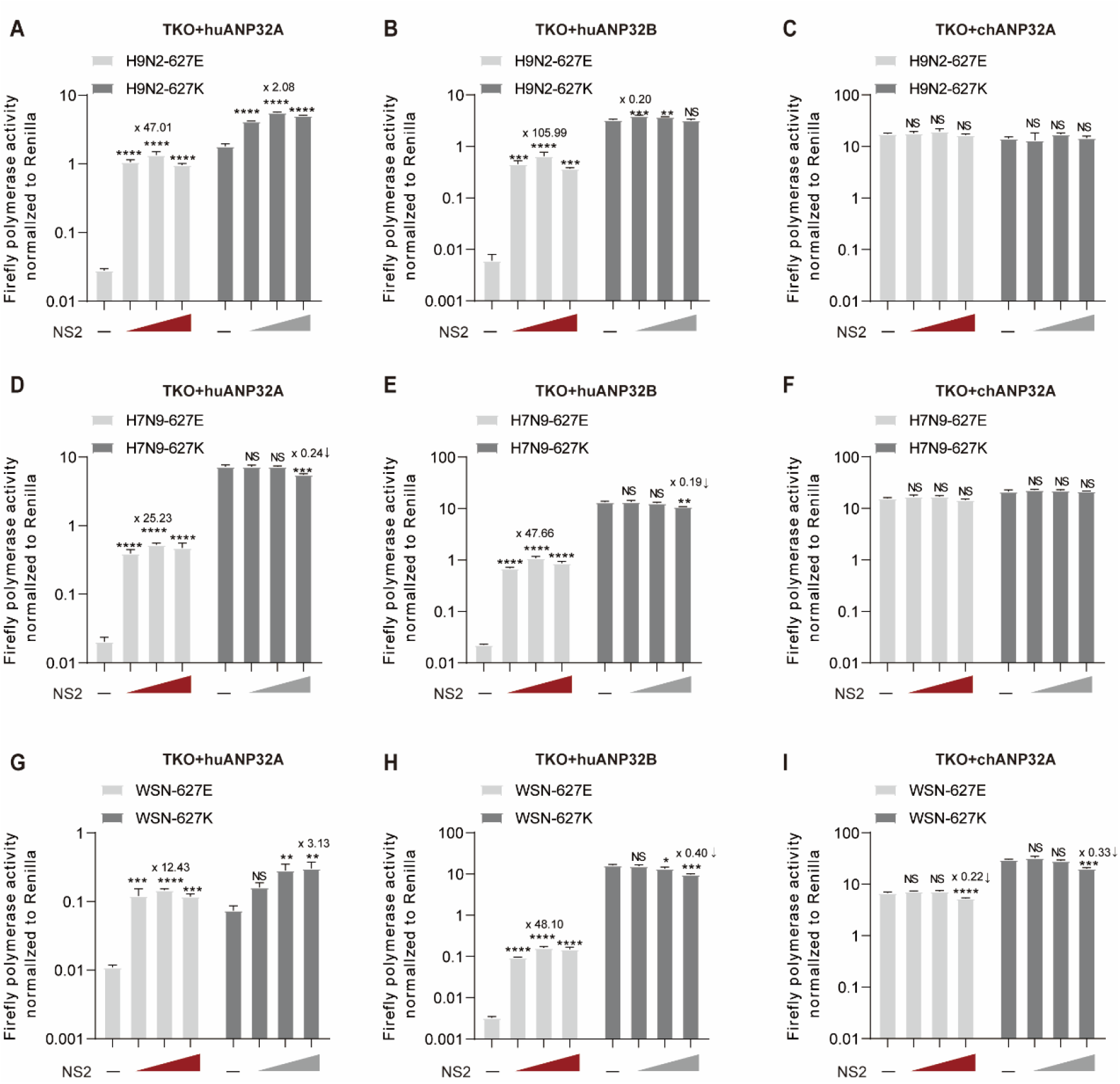
NS2 selectively enhances avian-signature vPol activity when supported by huANP32/B but not by chANP32A. **A-C:** vPol reconstitution assay to compare the effect of H9N2-NS2 on the activities of H9N2 PB2-627E and H9N2 PB2-627K vPols in TKO cells reconstituted with huANP32A (A), huANP32B (B) or chANP32A(C). **D-F:** Similar to A-C, but using NS2 and RNP proteins from avian influenza H7N9_AH13_. **G-I:** Similar to A-C, but using NS2 and RNP proteins from the A/WSN/33 virus. In (A) – (I), bars represent mean values of the replicates within one representative experiment (n = 3, ±SD). Vec, empty vector control. Significance was determi ned using one-way ANOVA followed by a Dunnett’s multiple comparisons test (\**p* < 0.05; \*\**p* < 0.01; \*\*\**p* < 0.001; \*\*\*\**p* < 0.0001; NS, non-significant).

To further investigate the general role of NS2 in enhancing huANP32A/B-mediated avian vPol activity, we used a human influenza H1N1_WSN_ polymerase to investigate the effect of H1N1_WSN_-NS2 on vPol activity of avian signature (PB2^627E^, WSN-627E) when supported by different ANP32 proteins. We found that H1N1_WSN_-NS2 also selectively enhanced the WSN-627E vPol activity when supported by huANP32A (∼12.43-fold increase) or huANP32B (∼48.10-fold increase), but not by chANP32A. H1N1_WSN_-NS2 also had a limited effect on the mammalian-signature (PB2^627K^, WSN-627K) vPol activity when supported by huANP32A, huANP32B or chANP32A (Figures 2G-I). These data suggest that the role of NS2 in selectively promoting the avian-signature vPol activity when supported by human ANP32A/B may not be virus strain-specific. To obtain additional evidence for this, we performed vPol reconstitution assays in TKO cells to test whether the NS2 proteins from IAVs isolated from different host species have a similar role in promoting avian-signature vPol activity when supported by huANP32A/B. As shown in Figures S1A-E, all NS2s of IAVs isolated from different taxa (including human, bird, pig, dog, and equine) have the ability to promote avian H9N2 and H7N9 (PB2^627E^) vPol activity when supported by huANP32A/B. However, the NS2 of influenza B virus (IBV) did not have this function.

The main known function of NS2 is to mediate the nuclear export of vRNP, which requires the nuclear export signal (NES) of NS2 and its binding to M1(Akarsu et al., 2003; Neumann et al., 2000; O’Neill et al., 1998; Paterson and Fodor, 2012). To test whether the avian-signature polymerase-enhancing properties of NS2 depend on its role in vRNP nuclear export, we generated a panel of NS2 mutants (Figure S2A), including those lacking NES or harboring a W78S point mutation that impairs the NS2-M1 interaction. The performance of these NS2 mutants was then evaluated in a vPol reconstitution assay in TKO cells. These NS2 mutants had the same positive effect on avian H9N2_ZJ12_ vPol activity when supported by huANP32A&B as did wild-type NS2 (Figure S2B), suggesting that the avian-signature polymerase-enhancing properties of NS2 is independent of its role in the mediation of vRNP nuclear export.

Together, these results suggest that influenza A virus NS2 can overcome the limitations of avian-signature vPol in human cells due to species-specific differences in the ANP32 protein.

### NS2 promotes avian-signature vPol activity when supported by ANP32A/B from different mammalian species

Previous studies have confirmed that ANP32A/B are critical factors for the function of influenza A virus polymerases from different species(Long et al., 2016; Long et al., 2019a; Staller et al., 2019; Zhang et al., 2019). Mammalian ANP32A/B proteins, including canine ANP32A/B (caANP32A/B), equine ANP32A/B (eqANP32A/B), mouse ANP32B (muANP32B), and swine ANP32B (swANP32B), lack a 33-residue insertion present in chANP32A, meaning that avian-signature vPol activity is restricted in most mammalian cells(Long et al., 2016; Peacock et al., 2020; Zhang et al., 2020a; Zhang et al., 2019). However, swANP32A contains a unique sequence feature that enables it to support avian vPol activity to a higher degree than other mammalian ANP32A/B(Peacock et al., 2020; Zhang et al., 2020a). Additionally, mouse ANP32A is inactive in supporting IAV polymerase activity, and only muANP32B supports IAV replication in mice(Staller et al., 2019; Zhang et al., 2019).

To examine the effect of NS2 on the activity of avian vPol when supported by caANP32A/B, eqANP32A/B, muANP32B or swANP32A/B, we performed a vPol reconstitution assay in TKO cells reconstituted with those ANP32A&B proteins. As shown in Figures 3A-D, H9N2-NS2 dramatically enhanced the activity of the H9N2 virus vPol when supported by the above-mentioned mammalian ANP32A/B, but had a relatively limited effect on the activity of the mammalian-signature (PB2^627K^) vPol. We obtained consistent results using NS2 and vPol from the avian H7N9_AH13_ virus (Figures 3E-H). These data suggest that NS2 can compensate for the defects of avian-signature vPol in mammalian cells, making it an indispensable cofactor in the adaptation of avian-signature polymerase to mammalian hosts.

**Figure 3.**
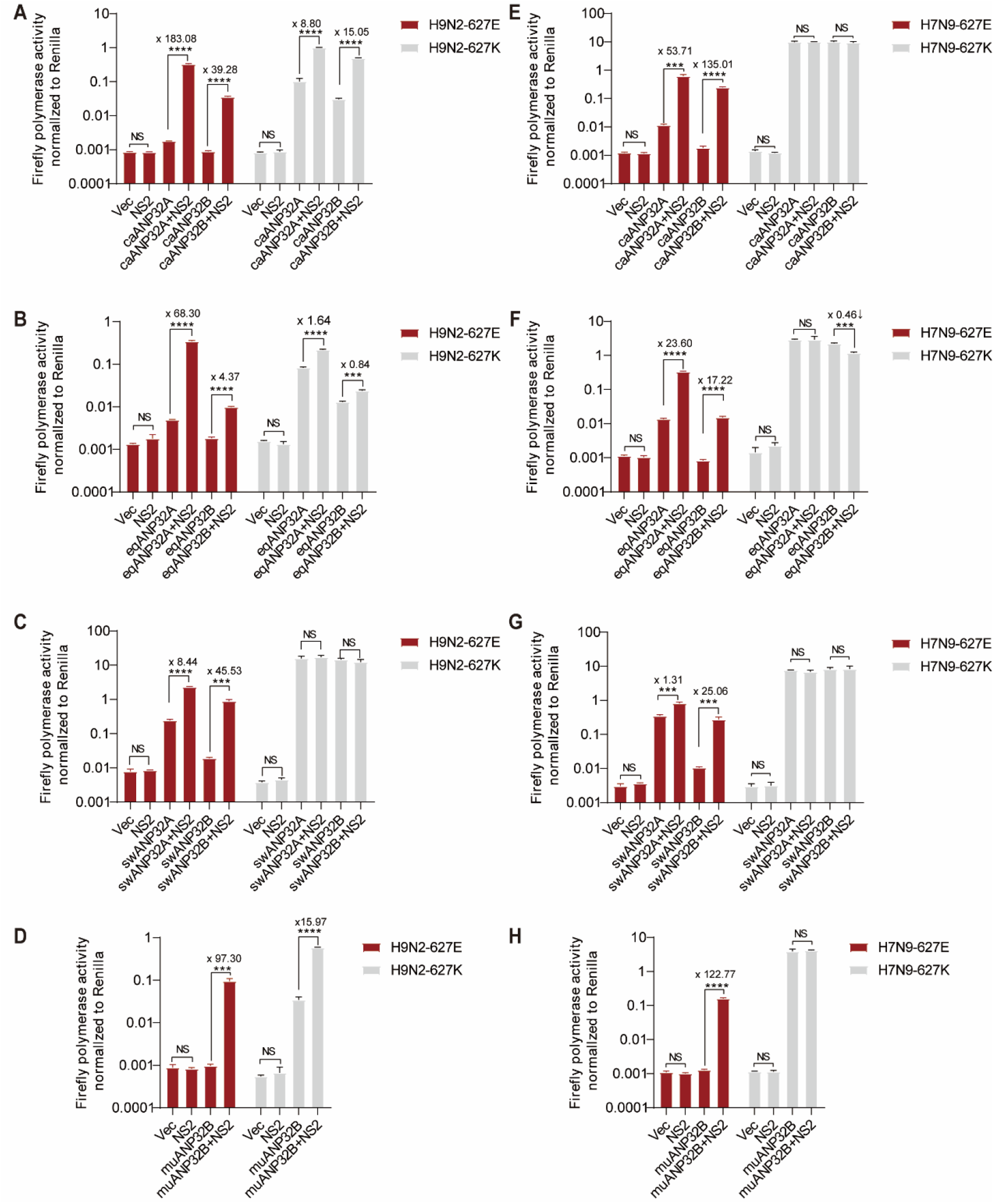
NS2 enhances avian-signature vPol activity when supported by ANP32A/B from different mammalian species. **A-D:** vPol reconstitution assay comparing the effect of H9N2-NS2 on H9N2 PB2-627E and H9N2 PB2-627K vPols activities in TKO cells reconstituted with ANP32A/B from dog (A), equine (B), pig (C) and mouse (D). **E-H:** Similar to A-D, but using NS2 and RNP proteins from avian influenza H7N9_AH13_. In (A)–(H), bars represent mean values of the replicates within one representative experiment (n = 3, ±SD). Vec, empty vector control. Significance was determined using an unpaired Student’s *t*-test (\*\*\**p* < 0.001; \*\*\*\**p* < 0.0001; NS, non-significant).

### The molecular mechanism by which NS2 selectively promotes avian-signature vPol activity in mammalian cells

Upon nuclear import, viral polymerase subunits (PA/PB1/PB2) assemble into the trimeric polymerase complex, which, together with genomic RNA and NP, further assemble into vRNPs(Eisfeld et al., 2015; Resa-Infante et al., 2011; Wandzik et al., 2021). Previous reports have shown that assembly of the trimeric viral polymerase is unaffected by the identity of amino acid 627 in the PB2 (K627 vs E627)(Mehle and Doudna, 2008). In contrast, vRNP assembly defects are associated with the restriction of avian-signature vPol activity in mammalian cells(Baker *et al*., 2018; Labadie et al., 2007; Mehle and Doudna, 2008). To gain insight into the mechanism by which NS2 enhances avian-signature vPol activity in mammalian cells, we first confirmed these observations by monitoring vRNP assembly in TKO cells reconstituted with huANP32A/B or chANP32A. In the presence of huANP32A/B, the amount of avian-signature polymerase (PB2-627E and PA) that co-precipitated with NP was less than that of mammalian-signature polymerase (PB2-627K and PA). However, in the presence of chANP32A, the co-precipitation of avian-signature polymerase (PB2-627E and PA) was increased (Figure 4A). This species-specific effect of ANP32 protein on avian signature vRNP assembly may determine the restriction of avian-signature vPol activity in mammalian cells.

**Figure 4.**
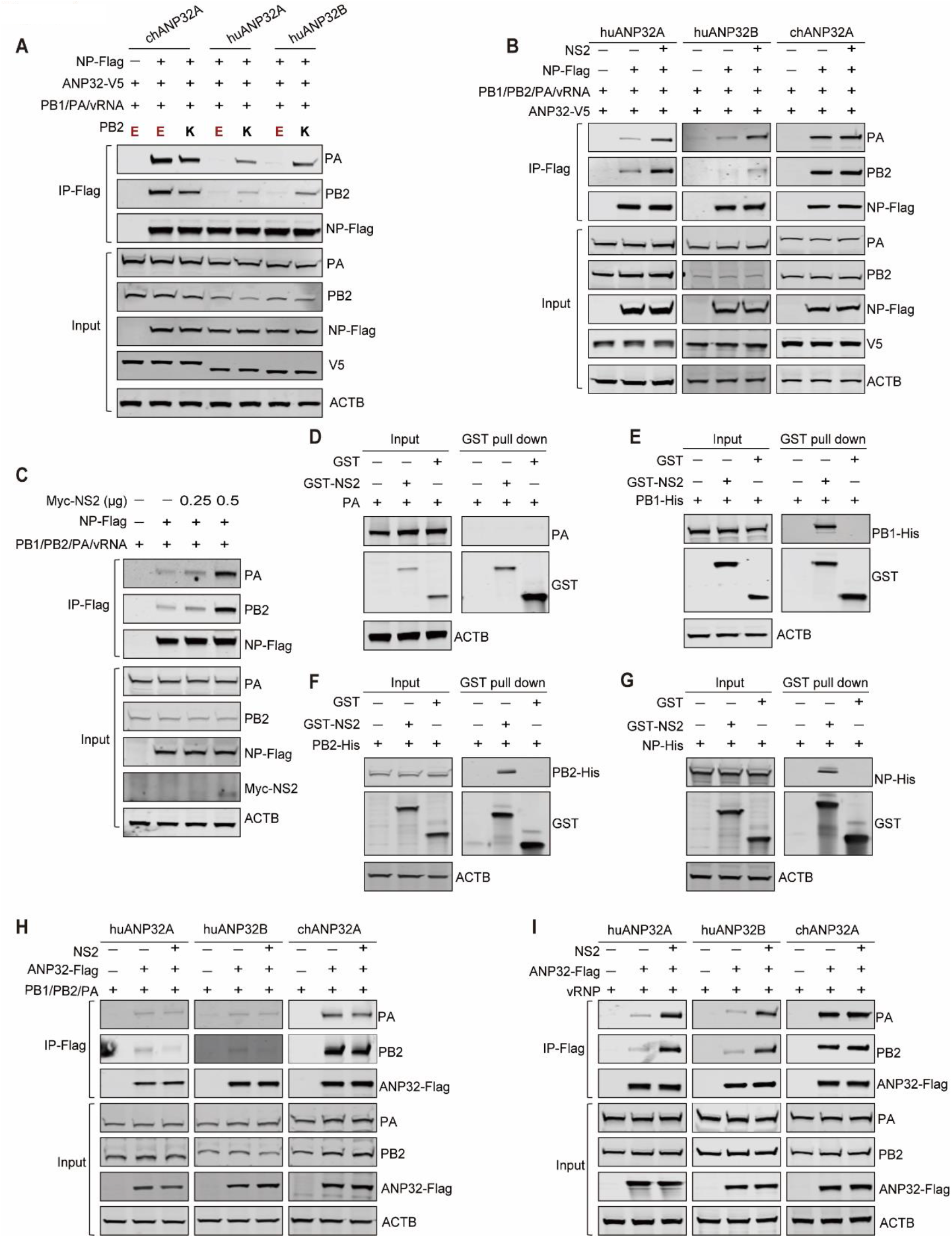
NS2 promotes the avian vRNP assembly and its interaction with human ANP32A/B, but not chANP32A. **A:** Comparison of avian (H9N2-PB2-627E) and mammalian-signature (H9N2-PB2-627K) vRNP assembly in TKO cells reconstituted with huANP32A/B or chANP32A. TKO cells were transfected with different V5-tagged ANP32A (0.4 μg), Flag-tagged NP (0.8 μg) and polymerase plasmids from avian influenza viruses H9N2_ZJ12_ (0.2 μg PA, 0.4 μg PB1, and 0.4 μg WT or E627K PB2) together with vRNA-luciferase reporter (0.4 μg). After anti-Flag precipitation at 24 h post-transfection, the indicated proteins were analyzed by western blotting. **B:** Comparison of the effect of NS2 on the avian vRNP assembly in TKO cells reconstituted with either huANP32A/B or chANP32A. TKO cells were transfected with different V5-tagged ANP32A (0.4 μg), Flag-tagged NP (0.8μg) and polymerase plasmids from avian influenza viruses H9N2_ZJ12_ (0.2 μg PA, 0.4 μg PB1, and 0.4 μg PB2) together with vRNA-luciferase reporter (0.4 μg) and Myc-tagged H9N2_ZJ12_-NS2 (50 ng). After anti-Flag precipitation at 24 h post-transfection, the indicated proteins were analyzed by western blotting. **C:** Comparison of the effect of NS2 on the avian-signature vRNP assembly in TKO cells without reconstitution of ANP32 proteins. TKO cells were transfected with Flag-tagged NP (0.8 μg) and polymerase plasmids from avian influenza viruses H9N2_ZJ12_ (0.2 μg PA, 0.4 μg PB1, and 0.4 μg PB2) together with vRNA-luciferase reporter (0.4 μg) and different doses of H9N2_ZJ12_-NS2 (0.25 μg, 0.5 μg). Following anti-Flag precipitation at 24 h post-transfection, the indicated proteins were analyzed by western blotting. **D-G:** GST affinity-isolation analysis of the complex formation of NS2 with either PA (D), PB1 (E), PB2 (F) or NP (G). GST pull-down assays were performed with lysates from HEK293T cells transfected with GST-tagged constructs (1 μg), as well as the indicated polymerase subunits (1 μg) and NP (1 μg) of avian influenza H9N2_ZJ12_ virus. **H:** Effect of NS2 on the binding of ANP32 proteins to avian-signature polymerase trimeric complexes. TKO cells were transfected with the different Flag-tagged ANP32 constructs (0.4 μg), together with Myc-tagged H9N2_ZJ12_-NS2 (50 ng) and polymerase plasmids (0.2 μg PA, 0.4 μg PB1, and 0.4 μg PB2) from the avian influenza H9N2_ZJ12_ virus. Following anti-Flag precipitation at 24 h post-transfection, the indicated proteins were analyzed by western blotting. **I:** Effect of H9N2-NS2 on the interaction between ANP32 proteins and avian-signature vRNP. Experiments were performed as in Fig.4H, except that the NP (0.8 μg) and vRNA-luciferase reporter (0.4 μg) were included in the transfection.

Next, to confirm whether the enhanced avian-signature vPol activity conferred by NS2 results from enhanced binding of the viral polymerase to the NP in mammalian cells, we assessed the effect of NS2 on the vRNP assembly in TKO cells reconstituted with either huANP32A/B or chANP32A. Immunoprecipitation assays showed that when TKO cells were reconstituted with huANP32A/B, the amounts of the H9N2_ZJ12_ polymerase subunits PA and PB2 co-precipitated by NP increased in the presence of NS2, whereas when chANP2A was present, co-precipitation of PB2 and PA by NP was not affected by NS2 (Figure 4B). These results confirm that the increased avian-signature vPol activity conferred by NS2 correlates with increased vRNP assembly in mammalian cells.

To further investigate the role of NS2 in facilitating vRNA assembly, the assembly process of H9N2 vRNP was monitored in TKO cells without reconstitution of any ANP32 proteins. In this case, vPol functions poorly during replication and transcription, resulting in a stable level of vRNA. However, to our surprise, NS2 was still able to promote the assembly of avian-signature vRNP (Figure 4C). In addition, we used GST pull-down analysis to demonstrate that NS2 associates with H9N2 RNP proteins, including PB1, PB2 and NP, but not PA (Figures 4D-G). These data strongly suggest that, like chANP32A, NS2 can compensate for the defective assembly of avian vRNP in mammalian cells and rescue avian-signature vPol activity in these cells through its association with RNP proteins.

Although there are many unknowns regarding the specific molecular mechanism by which the ANP32 protein supports the vPol activity of the influenza A virus, current evidence suggests that the interaction of ANP32 with the trimeric polymerase complex is critical for its function in supporting polymerase activity(Baker *et al*., 2018; Domingues and Hale, 2017; Long *et al*., 2019a; Staller *et al*., 2019; Sugiyama *et al*., 2015; Zhang *et al*., 2019). In addition, recent studies have shown that a unique feature (106V/156S) of swine ANP32A allows the protein to bind avian-signature trimeric polymerase complexes more strongly than other mammalian ANP32 proteins, and enabling it to support avian-signature vPol activity to a greater extent(Peacock *et al*., 2020; Zhang *et al*., 2020a). Therefore, we propose that NS2 promotes the avian-signature vPol activity in mammalian cells by enhancing the interaction between mammalian ANP32 proteins and the trimeric polymerase complex. To our surprise, H9N2-NS2 did not affect the interaction between huANP32A/B and the trimeric H9N2 virus polymerase complex, and neither was the interaction between chANP32A and the trimeric polymerase complex affected by NS2 (Figure 4H). However, when we explored whether NS2 affected the interaction between huANP32A/B and the H9N2 vRNP, we found that the amount of PA and PB2 co-precipitated by huANP32A/B was increased in the presence of NS2. NS2 did not affect the interaction between chANP32A and the vRNP (Figure 4I). Together, these data suggest that the rescue of avian-signature vPol activity by NS2 in mammalian cells correlates with its ability to promote the assembly process of avian vRNP and mammalian ANP32A/B-vRNP interactions.

### The SUMO interacting motif of NS2 determines its avian-signature polymerase-enhancing properties

Unlike mammalian ANP32A/B, chANP32A contains a 33-residue insertion, which allows it to effectively support the avian vPol activity(Long *et al*., 2016). Further evidence suggests that the presence of a SUMO interaction motif (SIM)-like sequence in this 33-residue insertion is critical for this supporting function(Domingues and Hale, 2017), suggesting that a SUMO-dependent function plays an important role in chANP32A-mediated avian vPol function. Therefore, we hypothesized that the SUMO-dependent function of NS2 might be related to its ability to promote avian-signature vPol activity in mammalian cells. To our surprise, when we used the in silico prediction program GPS-SUMO(Zhao et al., 2014), we found that H9N2-NS2 contains a SUMO-interacting motif (SIM) (^106^LLEVE^110^) (Figure 5A). A comparison of the amino acid sequence of this region (aa 106 to 110) with those from NS2 proteins from other viruses revealed that this SIM was highly conserved among influenza A viruses isolated from different host species, but not influenza B virus (Figure 5B). In addition, the crystal structure of the NS2 C-terminal domain(Akarsu *et al*., 2003) showed that the SIM was located on the C2 helix (Figure 5C). The presence of a SIM in NS2 suggested that this protein can interact non-covalently with SUMO1/2/3 or SUMO1/2/3-conjugated proteins. To gain more evidence for this, we performed SUMO binding assays in HEK293T cells by co-transfecting GST-tagged NS2 with His-Ubc9 and Flag-SUMO1/2/3. As shown in Figure 5D, GST-NS2, but not GST, could be co-precipitated by Flag-tagged SUMO1/2/3. However, the mutants defective in SUMO conjugation (SUMOm) failed to precipitate NS2, suggesting that NS2 specifically interacts non-covalently with SUMO1/2/3 conjugated proteins. Furthermore, the deletion of this SIM resulted in a failure of NS2 binding to Flag-SUMO1 conjugated proteins, further confirming that LLEVE is a bona fide SIM (Figure 5E).

**Figure 5.**
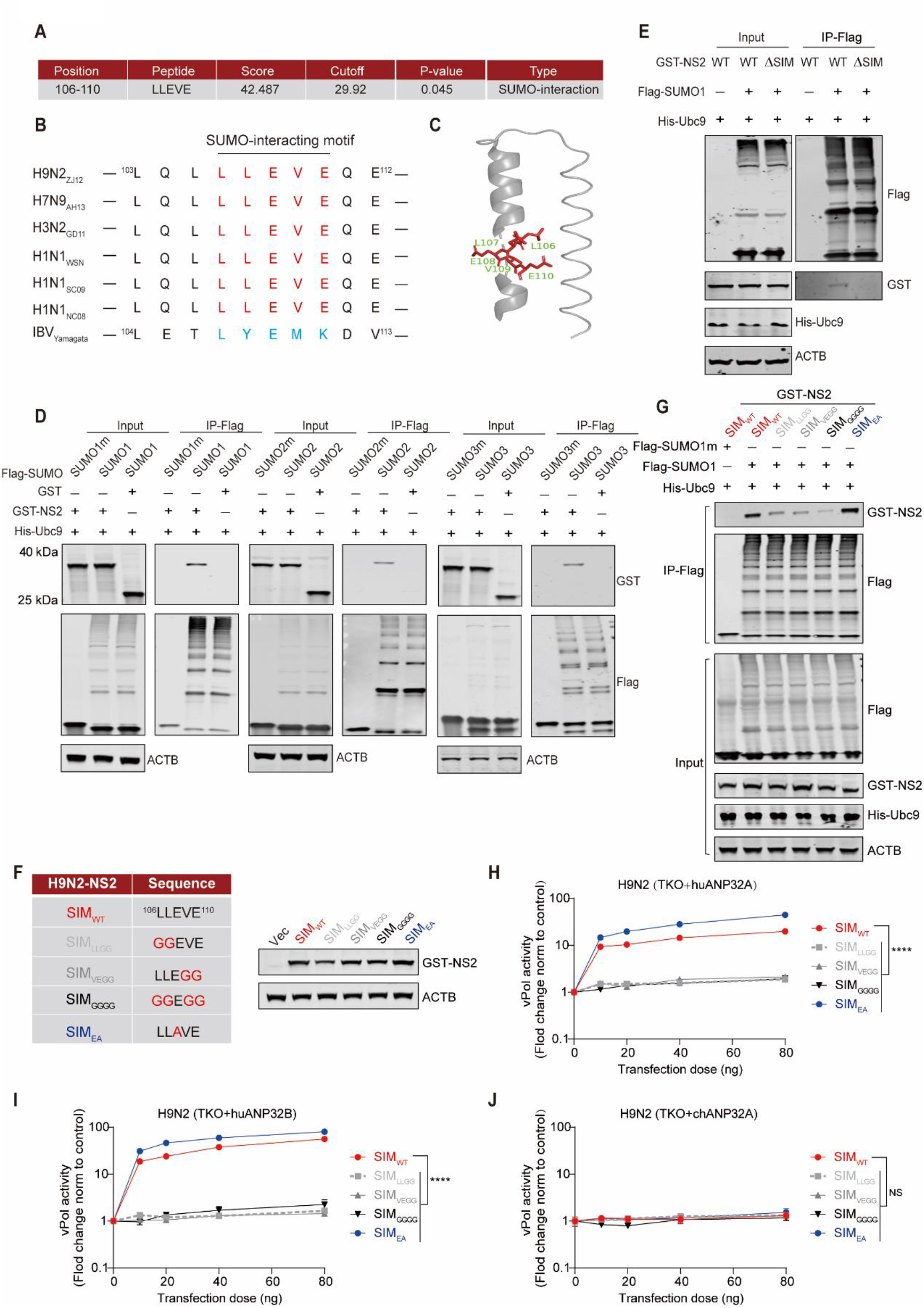
The SIM in NS2 determines its avian-signature polymerase-enhancing function. **A:** The candidate SUMO-interacting motif (SIM) in H9N2-NS2 was predicted using the GPS-SUMO software. **B:** Sequence alignment of NS2-SIM of influenza A viruses isolated from different host species and influenza B virus. **C:** Position of SIM in the structure of the NS2 C-terminal domain. **D:** In vivo SUMO binding assays. HEK293T cells were transfected with plasmids expressing GST-tagged NS2 from H9N2_ZJ12_ (1 μg), together with 1 μg Flag-tagged SUMO1/SUMO1m (left), SUMO2/SUMO2m (middle) and SUMO3/SUMO3m (right) and His-tagged Ubc9 (1 μg). At 24 h post-transfection, cell lysates were prepared for anti-Flag precipitation and the indicated proteins were analyzed by western blotting. **E:** In vivo SUMO binding assays. HEK293T cells were transfected with plasmids expressing GST-tagged NS2 or GST-tagged NS2 lacking SIM (ΔSIM) from H9N2_ZJ12_ (1 μg), together with 1μg Flag-tagged SUMO1 and His-tagged Ubc9 (1 μg). At 24 h post-transfection, cell lysates were prepared for anti-Flag precipitation and the indicated proteins were analyzed by western blot. **F:** A panel of generated NS2 mutants. The accompanying western blots show the effect of NS2 SIM changes on protein expression levels in these mutants. HEK293T cells were transfected with equal amount of indicted GST-tagged NS2 constructs. At 24 h post-transfection, cell lysates were prepared with RIPA lysis buffer and subjected to SDS-PAGE, followed by analysis with western blotting. **G:** The effects of NS2 SIM mutations on NS2 binding to Flag-tagged SUMO1. Experiments were performed as in Fig.5E. **H-J:** vPol reconstitution assay to compare the effect of NS2 SIM mutations on H9N2 vPol activity in TKO cells reconstituted with huANP32A (H), huANP32B (I) and chANP32A (J). TKO cells were transfected with different ANP32 constructs (20 ng), and polymerase plasmids (20 ng PA, 40 ng PB1 and 40 ng PB2) from the avian influenza H9N2_ZJ12_ virus together with vRNA-luciferase reporter (40 ng), 80 ng NP and increasing dose of H9N2_ZJ12_-NS2 plasmids (0-80 ng). For all assays, data were firefly activity normalized to *Renilla*, plotted as Fold change to empty vector (0 ng NS2 constructs). In (H)–(J), bars represent mean values of the replicates within one representative experiment (n = 3, ± SD). Vec, empty vector control. Significance was determined by two -way ANOVA (\*\*\*\**p* < 0.0001; NS, non-significant).

Functional SIMs are characterized by an arrangement of hydrophobic residues, and replacement of these hydrophobic residues with glycine impairs SIM-mediated function, as evidenced in previous reports(Domingues and Hale, 2017). The same approach was used here to generate a set of NS2 mutants to explore the contribution of the SIM in NS2 towards its avian polymerase-enhancing function. The NS2 protein expression level was not dramatically affected by these amino acid changes in NS2-SIM (Figure 5F). As expected, substitution of these hydrophobic residues in this motif for glycines (SIM_LLGG_, SIM_VEGG_ and SIM_GGGG_) severely impairs the interaction between Flag-SUMO1-conjugated proteins and NS2, while substitution at the non-hydrophobic position E108 (SIM_EA_) in this SIM did not (Figure 5G). Next, the effects of these NS2 mutants on avian vRNP assembly and avian vRNP-ANP32A interactions were assessed in TKO cells reconstituted with either huANP32A or chANP2A. Surprisingly, in TKO cells reconstituted with huANP32A, NS2 mutants in which hydrophobic residues in the SIM were replaced with glycines lost the ability to promote avian vRNP assembly, while SIM_EA_ retained this ability (Figure S3A). In contrast, wild type and mutant NS2 did not affect avian vRNP assembly in the presence of chANP32A (Figure S3A). Avian vRNP-huANP32A interactions were enhanced in the presence of wild type NS2 and SIM_EA_, but not in NS2 mutants with glycine substitutions in the SIM (Figure S3B). None of the NS2 constructs affected the avian vRNP-chANP32A interaction (Figure S3B).

Subsequent vPol reconstitution assays were performed in TKO cells, to assess the effect of NS2 mutants on avian-signature vPol activity in TKO cells reconstituted with huANP32A, huANP32B or chANP32A. As shown in Figures 5H and 5I, the NS2-SIM_EA_ mutant was even more potent than WT-NS2 in promoting avian H9N2_ZJ12_ vPol activity supported by huANP32A/B, whereas the NS2-SIM_LLGG_, NS2-SIM_VEGG_ and NS2-SIM_GGGG_ mutants almost completely lost this ability. Notably, both wild-type NS2 and these mutants had only a limited effect on the avian H9N2_ZJ12_ vPol activity supported by chANP32A (Figure 5J). We confirmed these observations using NS2 and RNP proteins from an H7N9 AIV genetic background. Similarly, H7N9-NS2 mutants in which the hydrophobic residues in the SIM were substituted with glycines lost the avian-signature polymerase-enhancing properties, while NS2-SIM_EA_ did not (Figures S4A-D). None of the H7N9-NS2 mutants had more than a limited effect on H7N9 (PB2-627E) vPol activity when supported by chANP32A (Figure S4D).

The above data indicate that the SIM in NS2 is required for its avian-signature polymerase-enhancing function. Therefore, we next assessed the prevalence of this motif among influenza A viruses. The SIM (^106^LLEVE^110^) was highly conserved in avian isolates of H1-H16 influenza viruses (Table 1). In addition, we focused on several subtypes of avian influenza viruses capable of cross-species transmission, including H5N1, H7N9, H5N6, H5N8 and H9N2. The analysis showed that their NS2 SIMs were similarly well conserved (Table 1).

Previous studies have shown that NS2 can be SUMOylated(Domingues et al., 2015; Pal et al., 2011). To assess the contribution of the SUMOylation of NS2 to its avian-signature polymerase-enhancing properties, we generated an NS2 mutant with lysine-to arginine loss-of-function substitutions at all six lysine residues of the protein (termed NS2-K0) (Figure S5A). Surprisingly, NS2-K0 still retained the ability to promote the avian-signature vPol activity when supported by huANP32A/B (Figures S5B-C). However, like wild type NS2, NS2-K0 had only a very limited effect on chANP32A-mediated avian-signature vPol function (Figure S5D). Meanwhile, the NS2 mutant lacking SIM (NS2-ΔSIM) lost its avian-signature polymerase-enhancing properties (Figures S5B-D). These data suggest that NS2 selectively relies on its SIM to exert its avian-signature polymerase–enhancing function.

### SIM of NS2 is crucial for efficient replication of avian influenza virus in mammalian cells

Our vPol reconstitution assays identified NS2 as a cofactor necessary for the adaptation of avian-signature polymerase to mammalian hosts. To determine whether the species-specific effect on the avian-signature vPol activity conferred by NS2 manifested as species-specific changes in the replication of avian influenza virus in mammalian cells, we first intended to rescue H9N2 avian influenza viruses carrying different mutant SIMs on NS2, which cause defects in promoting avian-signature polymerase activity in mammalian cells. Unfortunately, we were unable to rescue the mutant avian influenza H9N2 viruses bearing NS2 SIM_LLGG_, SIM_VEGG_ or SIM_GGGG_ mutations, which have almost completely lost their avian-signature polymerase-enhancing properties. However, the mutant avian influenza H9N2 virus with NS2 bearing the SIM_E110G_ mutation, which partially impaired its ability to bind SUMO1-conjugated proteins and promote avian-signature vPol activity supported by huANP32A/B (Figures S6A-C), was successfully rescued. We next performed multicycle replication assays to compare the replication ability of wild-type (SIM_WT_) and mutant (SIM_E110G_) avian influenza virus H9N2 in different cell lines (Figures 6A-D). The replication kinetics and titer of the H9N2-SIM_E110G_ virus in primary chicken embryo fibroblasts were comparable to those of the H9N2-SIM_WT_ virus (Figure 6A), which is consistent with the fact that chANP32A mediated avian-signature vPol function was not greatly affected by the NS2. However, replication of the H9N2-SIM_E110G_ virus was significantly impaired in A549 and MDCK cells (Figures 6B-C). Notably, the replication defect of H9N2-SIM_E110G_ virus in MDCK cells was completely rescued upon overexpression of chANP32A in these cells (Figure 6D). These data suggest that the species-specific effect on the avian-signature vPol activity conferred by NS2 also affects the replication of the avian influenza virus in a species-specific fashion. Consistent results were obtained when multicycle replication assays were performed using the two highly pathogenic avian influenza viruses H7N9_HN17_ (Figures 6E-H) and H5N6_GD16_ (Figures 6I-L). In addition, to obtain more evidence to support the general role of NS2 in promoting avian influenza virus replication in mammalian cells, we used a recombinant A/WSN/33 virus encoding PB2 K627E (WSN-627E), which shows significant replication defects in mammalian cells due to host restrictions as previously described(Mehle and Doudna, 2008), as an avian-like virus in the following experiment. As we expected, the WSN-627E-SIM_E110G_ virus showed replication defects in A549 and MDCK cells (Figures 6N-O), but no change in viral titers was detected from primary chicken embryo fibroblasts (Figure 6M) and MDCK-chANP32A cells (Figure 6P). Together, these results confirm the essential role of the SIM of NS2 in promoting the adaptation of avian influenza virus to mammalian cells.

**Figure 6.**
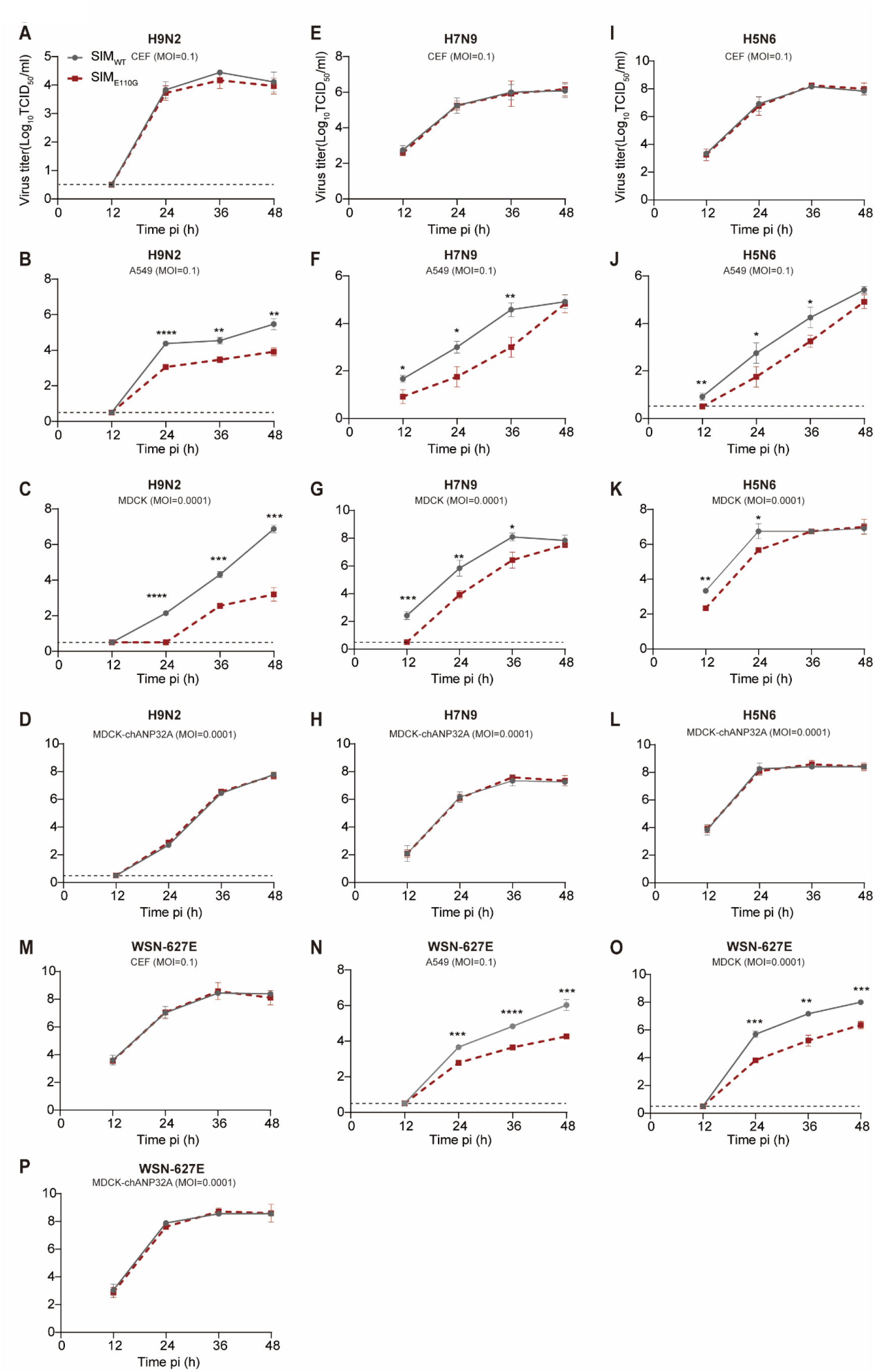
SIM of NS2 promotes avian influenza virus replication in mammalian cells but not in avian cells. **A-D:** Comparison of the replication abilities of the avian influenza virus H9N2_ZJ12_ (H9N2-SIM_WT_) and the avian influenza virus H9N2 with NS2 SIM harboring an E110G mutation (H9N2-SIM_E110G_) in indicated cell lines. Primary chicken embryonic fibroblasts (CEF) (A), A549 cells (B), MDCK cells (C) and MDCK-chANP32A cells (MDCK cells stably expressing chANP32A) (D) were infected with the H9N2-SIM_WT_ virus or the H9N2-SIM_E110G_ mutant. Cell supernatants were collected at the described time-points and virus titers were determined on MDCK cells stably-expressing chANP32A using TCID_50_ assays. **E-H:**Comparison of the replication abilities of avian influenza virus H7N9 (SIM_WT_) and the avian influenza virus H7N9 with NS2 SIM harboring an E110G mutation (SIM_E110G_) in the indicated cell lines. Experiments were performed as in Fig.6A-D. **I-L:**Comparison of the replication abilities of the avian influenza virus H5N6 (H5N6-SIM_WT_) and the avian influenza virus H5N6 with NS2 SIM harboring an E110G mutation (H5N6-SIM_E110G_) in the indicated cell lines. Experiments were performed as in Fig.6A-D. **M-P:** Comparison of the replication abilities of the avian-signature WSN-627E virus (SIM_WT_) and the avian-signature WSN-627E virus with NS2 SIM harboring an E110G mutation (SIM_E110G_) in the indicated cell lines. Experiments were performed as in Fig.6A-D. In (A)–(P), bars represent mean values of the replicates within one representative experiment (n = 3. ± SD). Significance was determined at each time point using an unpaired Student’s t-test. Asterisks indicate significant differences between SIMWT and SIME110G (\**p* < 0.05; \*\**p* < 0.01; \*\*\**p* < 0.001; \*\*\*\**p* < 0.0001).

### The replication and virulence of the avian influenza viruses H5N6 with NS2-SIM_E110G_ utations are attenuated in mice, but not in chickens

To determine the importance of avian-signature polymerase-enhancing properties of NS2 on the replication capacity and virulence of avian influenza virus in mice, the H5N6_GD16_ avian influenza virus was selected for the following in vivo experiments. H5N6_GD16_ was selected because it has been widely detected in wild birds and poultry, and because in recent years it has also been reported to infect humans, posing a huge threat, and causing high mortality in human populations(Bi et al., 2016). To compare the replication ability of the wild type H5N6_GD16_ (H5N6-SIM_WT_) virus and the mutant H5N6_GD16_ (H5N6-SIM_E110G_) virus in mice, two groups of five 6-week-old female BALB/c mice were inoculated with these viruses at an infectious dose of either 10^2^ EID_50_ or 10^4^ EID_50_, respectively. The nasal turbinates and lungs of the mice were collected on days 3, 5 and 7, and the viral titer was determined in MDCK-chANP32A cells using TCID_50_ assays. The replication ability of the H5N6-SIM_E110G_ virus in the lungs of mice was significantly reduced at an infectious doses of both 10^2^ EID_50_ (Figure 7B) and 10^4^ EID_50_ (Figure 7D) on day 5 p.i and day 7 p.i. At the low infectious dose, the replication ability of H5N6_GD16_ in nasal turbinate was relatively low, and under these conditions, the viral titers in the nasal turbinates of mice infected with the H5N6-SIM_E110G_ virus were similar to those of mice infected with the H5N6-SIM_WT_ virus (Figure 7A). However, when mice were inoculated with 10^4^ EID_50_ of the H5N6 virus, the viral titers in the nasal turbinates of those infected with the H5N6-SIM_E110G_ virus were lower (*p* = 0.0513 by *t* test) than of those infected with the H5N6-SIM_WT_ virus on day 5 p.i (Figure 7C). Groups of 6-week-old female BALB/c mice were then inoculated with serial dilutions of the H5N6-SIM_WT_ virus or the H5N6-SIM_E110G_ virus, and the MLD_50_ values of these viruses were determined and the subjects monitored for mortality for two weeks. As shown in Figure 7E, the MLD_50_ of the H5N6-SIM_WT_ virus was 3.22 Log_10_EID_50_, whereas the MLD_50_ of the H5N6-SIM_E110G_ virus was reduced approximately 19.05-fold (MLD50 = 4.50 Log_10_EID_50_). These results suggest that the replication and virulence of the H5N6 avian virus in mice also depend on the avian-signature polymerase-enhancing properties of the NS2 SIM.

**Figure 7.**
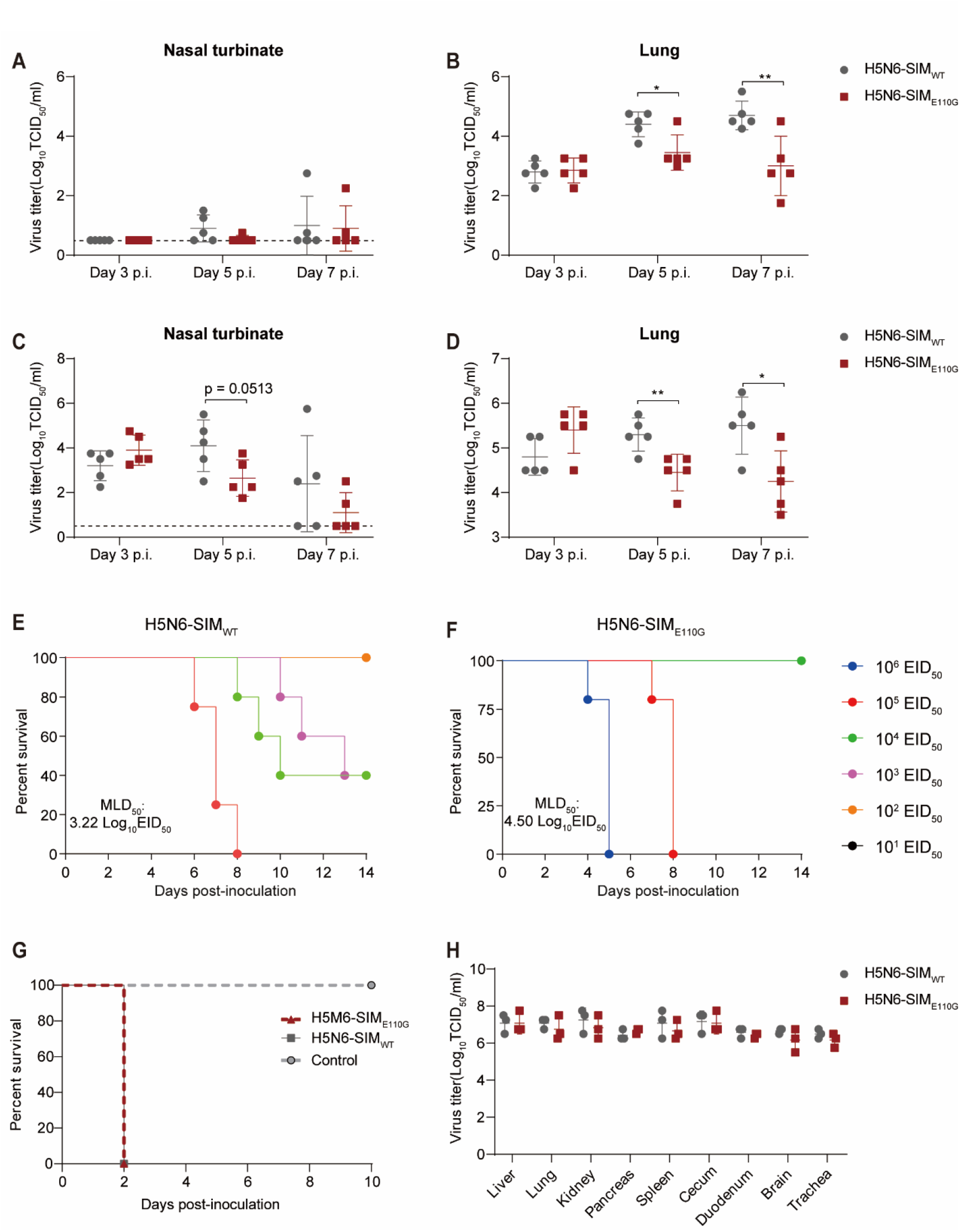
SIM of NS2 affects the replication and virulence of the avian influenza virus H5N6 in mice but not in chickens. **A-D:** Replication of the H5N6-SIM_WT_ virus and the H5N6-SIM_E110G_ mutant virus in mice. Five 6-week-old BALB/c mice were inoculated intra-nasally with the H5N6-SIM_WT_ virus or the H5N6-SIM_E110G_ mutant virus at an infectious dose of either 10^2^ EID_50_ (A-B) or 10^4^ EID_50_ (C-D), respectively. Nasal turbinates (A, C) and lungs (B, D) of mice were collected on day 3 p.i, 5 p.i and 7 p.i for virus titration in MDCK-chANP32A cells using TCID_50_ assays. **E-F:** MLD_50_ for mice infected with the H5N6-SIM_WT_ virus (E) or the H5N6-SIM_E110G_ virus (F). **G:** Mortality of chickens inoculated with the H5N6-SIM_WT_ virus or the H5N6-SIM_E110G_ virus. **H:** Viral titers in the organs of chickens inoculated with the H5N6-SIM_WT_ virus or the H5N6-SIM_E110G_ virus. Three chickens were inoculated i.n. with 10^6^ EID_50_ of the H5N6-SIM_WT_ virus or the H5N6-SIM_E110G_ mutant virus in a 0.1 ml volume. The organs, including brains, lungs, kidneys, spleens, pancreas, duodenums, liver, tracheae, and ceca, were collected on day 3 p.i for viral titration in MDCK-chANP32A cells using TCID_50_ assays. In (A)–(D) and (H), bars represent mean values of the replicates within one representative experiment (n ≥ 3, ± SD). Significance was determined per time point using an unpaired Stu dent’s *t-*test. Asterisks indicate significant differences between H5N6-SIM_WT_ and H5N6-SIM_E110G_ (* *p* < 0.05; ** *p* < 0.01).

Next, to determine whether the replication and pathogenicity of H5N6 avian influenza virus in chickens were affected by the NS2-SIM_E110G_ mutation, an intravenous pathogenicity index (IVPI) test was performed using the H5N6-SIM_WT_ and H5N6-SIM_E110G_ viruses according to the recommendations of the Office International Des Epizooties(Office International des Epizooties). Groups of ten 6-week-old SPF chickens were inoculated intravenously (i.v.) with 0.1 ml of a 1:10 dilution of bacterium-free allantoic fluid containing either H5N6-SIM_WT_ or H5N6-SIM_E110G_ virus and were observed for signs of disease or death for 10 days. All chickens inoculated with either the wild type or the mutant virus died within 48 hours post-inoculation, yielding an IVPI value of 3 (Figure 7G). To compare the abilities of the H5N6-SIM_WT_ and H5N6-SIM_E110G_ viruses to replicate in chickens, two groups of three 6-week-old SPF chickens were inoculated intranasally with 10^6^ EID_50_ of either H5N6-SIM_WT_ or H5N6-SIM_E110G_ virus in a 0.1 ml volume. The chickens were killed on day 3 p.i., and their organs, including brains, lungs, kidneys, spleens, pancreas, duodenums, liver, tracheae, and ceca were collected for viral titration using TCID_50_ assays in MDCK-chANP32A cells. Both WT and H5N6-SIM_E110G_ virus replicated systemically with comparable titers in chickens (Figure 7H). These results indicated that NS2 SIM function is not required for replication of avian IAV in chickens.

Taken together, these results suggest that the replication and virulence of the avian influenza virus in mice, but not in chickens, is also dependent on the help of the NS2 proteins with a functional SIM.

## DISCUSSION

Mammalian ANP32A/B efficiently support the activity of mammalian-signature polymerase, but do not support avian vPol due to the species-specific differences in ANP32 proteins(Baker *et al*., 2018; Bi *et al*., 2019; Domingues *et al*., 2019; Domingues and Hale, 2017; Long *et al*., 2016; Long *et al*., 2019a; Peacock *et al*., 2020; Staller *et al*., 2019; Zhang *et al*., 2020a; Zhang *et al*., 2019). Therefore, avian influenza virus requires adaptive mutations, such as PB2-E627K, to adapt to mammalian ANP32 proteins and establish productive replication when jumping from birds to mammals(Long *et al*., 2016; Long *et al*., 2019a; Zhang *et al*., 2019). However, in some cases, certain isolates of avian influenza virus, such as H7N9 and H5N1, infect humans to establish productive infections and can be fatal without acquiring previously identified adaptive mutations(Guo *et al*., 2021; Li *et al*., 2014b; Liang *et al*., 2019; Shi *et al*., 2018; Yin *et al*., 2021; Zhao et al., 2012). The molecular mechanisms by which avian influenza viruses adapt to mammals without prior adaptation are currently unknown. Furthermore, the molecular mechanisms underlying the early-stage replication process of non-adapted avian influenza viruses prior to the emergence of adaptive mutations also remain to be elucidated. In this study, we show that NS2 selectively enhances the activity of the avian-signature vPol when supported by mammalian ANP32A/B but not chANP32A, thereby eliminating the restriction of avian-signature polymerase in mammalian cells. This avian-signature polymerase-enhancing property of NS2 is exerted by promoting the assembly of avian vRNP and the binding of avian vRNP to mammalian ANP32A/B. Most importantly, the presence of a SUMO-interacting motif within the C-terminal tail of NS2 is required to confer its avian-signature polymerase-enhancing ability. Interestingly, NS2 has a relatively limited effect on the activity of mammalian-signature (PB2-627K) vPol supported by mammalian ANP32A/B or chANP32A. Thus, we have identified NS2 as a potential alternative strategy for the avian influenza virus to overcome the restriction of polymerase and thus better adapt when it crosses the species barrier from birds to mammals.

The main known function of NS2 is to mediate the nuclear export of vRNPs, a process that requires the combined participation of its nuclear export signal and M1(Akarsu *et al*., 2003; Neumann *et al*., 2000; O’Neill *et al*., 1998; Paterson and Fodor, 2012). In addition, NS2 regulates the transcription and replication process of the influenza A/WSN/33 virus(Robb *et al*., 2009), and is also required to for the production of small viral RNAs (svRNAs)(Perez et al., 2010). A previous study found that adaptative mutations in NS2 promoted the adaptation of the avian influenza virus H5N1 to human cells(Manz et al., 2012). Here we discovered a novel function of NS2 in species-specific support of avian influenza virus replication in mammals in vitro and in vivo. Our results indicate that NS2 from avian and mammalian influenza A viruses can selectively enhance avian-signature vPol activity when supported by mammalian ANP32A/B, but not by chANP32A. However, the mammalian-signature vPol activity supported by mammalian ANP32A/B was limited affected by the NS2. Furthermore, we demonstrate that the avian-signature polymerase-enhancing function of NS2 is independent of its role in the vRNP nuclear export by generating a panel of NS2 mutants with defects in mediating vRNP nuclear export.

Previous studies have shown that the restriction of avian-signature vPol activity is associated with impaired vRNP assembly in mammalian cells(Baker *et al*., 2018; Labadie *et al*., 2007; Mehle and Doudna, 2008). Recent evidence suggests that the species-specific differences in ANP33A determine the restriction of avian-signature vPol activity in mammalian cells(Long *et al*., 2016). The restriction of avian-signature vPol activity in human cells could be compensated for by expression of chANP32A via unknown mechanisms(Baker *et al*., 2018; Long *et al*., 2016), and it is unclear whether chANP32A functions to support avian-signature vPol activity by promoting vRNP assembly in mammalian cells. Here, we provide evidence that the defects of avian-signature vRNP assembly in human cells could be compensated for by expression of chANP32A or NS2. However, when TKO cells are reconstituted with chANP32A, the avian vRNP assembly process is unaffected by NS2 expression. These data suggest that NS2 promotes avian vRNP assembly in a species-specific manner. Based on our findings, we propose that NS2 mediates the enhancement of avian vPol activity through a mechanism common to chANP32A. To exclude the possibility that the failure of avian vRNP assembly in TKO cells reconstituted with huANP32A/B was due to a limiting amount of vRNA due to low avian vPol activity when supported by huANP32A/B in the absence of NS2, the avian vRNP assembly process was assessed in TKO cells without reconstitution of any ANP32 proteins. In this case, the avian polymerase was not functional and vRNA levels were stable. However, we still found that NS2 promoted the assembly of avian vRNP (Figure 4C). These results suggest that the defects in avian vRNP formation do not arise directly from intrinsic differences in vPol activity. Furthermore, in addition to polymerase directing genome replication and transcription in the form of vRNP(Fodor and Te Velthuis, 2019; Wandzik *et al*., 2021), genomic segments are also packaged into virions in the form of vRNP. Therefore, we hypothesize that NS2 likely promotes avian influenza virus replication in mammalian cells not only by enhancing avian polymerase activity, but also through other mechanisms remaining to be determined.

Previous studies have shown that the polymerase-supporting function of ANP32 is associated with its association with vPol(Baker *et al*., 2018; Domingues and Hale, 2017; Long *et al*., 2019a; Staller *et al*., 2019; Sugiyama *et al*., 2015; Zhang *et al*., 2019). However, avian-signature vPol activity is poorly supported by the mammalian ANP32A, a fact which cannot be explained by differences in the interaction of ANP32A with vPol, because the binding of ANP32A to vPol is independent of PB2-627 identity(Domingues and Hale, 2017). Here, we found that NS2 promoted the avian-signature vPol activity supported by mammalian ANP32A/B without affecting the interaction between mammalian ANP32A/B and vPol. In contrast, we found that NS2 significantly enhanced the interaction between mammalian ANP32 protein and avian-signature vRNP. The avian-signature vPol activity supported by chANP32A was affected by NS2 only to a limited extent, which corresponds to the fact that NS2 protein had only a limited effect on the chANP32A-vRNP interaction.

SUMOylation is an important post-translational modification in which SUMO molecules can be covalently attached to the lysine of the target protein(Geiss-Friedlander and Melchior, 2007). In mammals, only three SUMO isoforms, SUMO1, SUMO2 and SUMO3, are associated with SUMOylation. SUMOylation typically targets lysines located within the consensus motif (ҨKxD/E, where Ҩ represents hydrophobic amino acid and x represents any amino acid) for covalent modification(Rodriguez et al., 2001; Sampson et al., 2001). Additionally, SUMO and target proteins can interact non-covalently through SUMO interacting motifs (SIMs)(Hecker et al., 2006; Song et al., 2004). Current studies have shown that multiple proteins of the influenza A virus, including NS1, M1, NP, PB1 and PB2, can be modified by SUMOylation(Guo et al., 2022; Han et al., 2014; Li et al., 2021; Wang et al., 2022; Wu et al., 2011; Xu et al., 2011). In this study, we found for the first time that NS2 contains a conserved SIM sequence in its C-terminal tail. NS2 can non-covalently bind to SUMO1/2/3-conjugated proteins through this SIM. Disruption of the integrity of this SIM by replacing these hydrophobic residues in this motif for glycines (SIM_LLGG_, SIM_VEGG_ and SIM_GGGG_) resulted in the loss of binding ability of the NS2 protein to SUMO1-conjugated proteins. Moreover, NS2 mutants with impaired SIM integrity failed to enhance avian vRNP assembly and the binding of avian vRNP to mammalian ANP32A/B, thereby almost completely losing their avian-signature polymerase-enhancing properties. Although we were unable to rescue recombinant avian influenza viruses H9N2 with NS2 bearing SIM_LLGG_, SIM_VEGG_ or SIM_GGGG_ mutations, which have almost completely lost their avian-signature polymerase-enhancing properties. Single SIM-E110G mutation is already sufficient to affect the replication of avian influenza virus in a species-specific manner.

Previous studies have shown that the presence of SIM typically enhances the SUMOylation of its own proteins(Lin et al., 2006; Martinat et al., 2021). Here, we provided evidence that the SUMOylation of NS2 is not required for its avian-signature polymerase-enhancing ability by generating a lysine-free mutant of NS2, suggesting that the SIM-mediated function of NS2 is independent of its SUMOylation. Furthermore, the failure of SUMO-conjugation-deficient mutants to bind to NS2 suggests that NS2 may interact with specific SUMO-conjugated viral proteins or other host factors through its SIM to exert its polymerase-enhancing function, which remains to be elucidated in future studies.

The mechanism underlying the early stages of avian influenza virus replication in mammalian cells, prior to the emergence of adaptative mutations, have long remained unknown. In most cases, the emergence of adaptative mutations was still required for better adaptation. Here, we provide evidence that non-adapted avian vPol can use mammalian ANP32A/B with the help of NS2 to replicate productively in the early stages before the emergence of adaptive mutations. Notably, SIM (^106^LLEVE^110^) is highly conserved in avian isolates of H1-H16 influenza A viruses. Based on our findings, we hypothesize that the replication advantage conferred by the avian-signature polymerase-enhancing properties of NS2 may facilitate the adaptation process by accelerating the acquirement of adaptive mutations. Adaptative mutations in NS2 have been reported previously. The adaptive mutation M16I in H5N1 NS2 was shown to compensate for the replication defect of H5N1 in human cells(Manz *et al*., 2012). Considering the NS2s from influenza A viruses isolated from different host species are conserved in this avian-signature polymerase-enhancing function, we reason that the M16I mutation only positively regulates this function. Since we demonstrate that the SIM in NS2 determines this function, it is possible that M16I might enhance the binding of NS2 to SUMO-conjugated proteins, an area which requires further investigation.

To conclude, our study identified NS2 as a cofactor in the adaptation of the avian influenza virus to mammalian hosts. Most importantly, we uncovered a previously unknown mechanism by which avian influenza viruses use the SIM in NS2 to assist in crossing the species barrier from birds to mammals.

## ACKNOWLEDGMENTS

We thank Dr. Yoshihiro Kawaoka and Dr. Zejun Li for providing plasmids. This study was supported by grants from the National Natural Science Foundation of China (31521005, 32002275), and the Natural Science Foundation of Heilongjiang Province of China (YQ2020C021 and TD2022C006).

## AUTHOR CONTRIBUTIONS

X.J.W. conceived and supervised the study and revised the manuscript; L.K.S., H.L.C., and X.J.W. analyzed the data and designed the study. L.K.S. wrote the original draft; L.K.S., H.H.K., M.Y.Y., L.N., Y.X.Q., Y.Z. performed experiments; Z.Y.Z and H.L.Z. provided reagents and resources.

## Supplemental Figures

**Figure S1.**
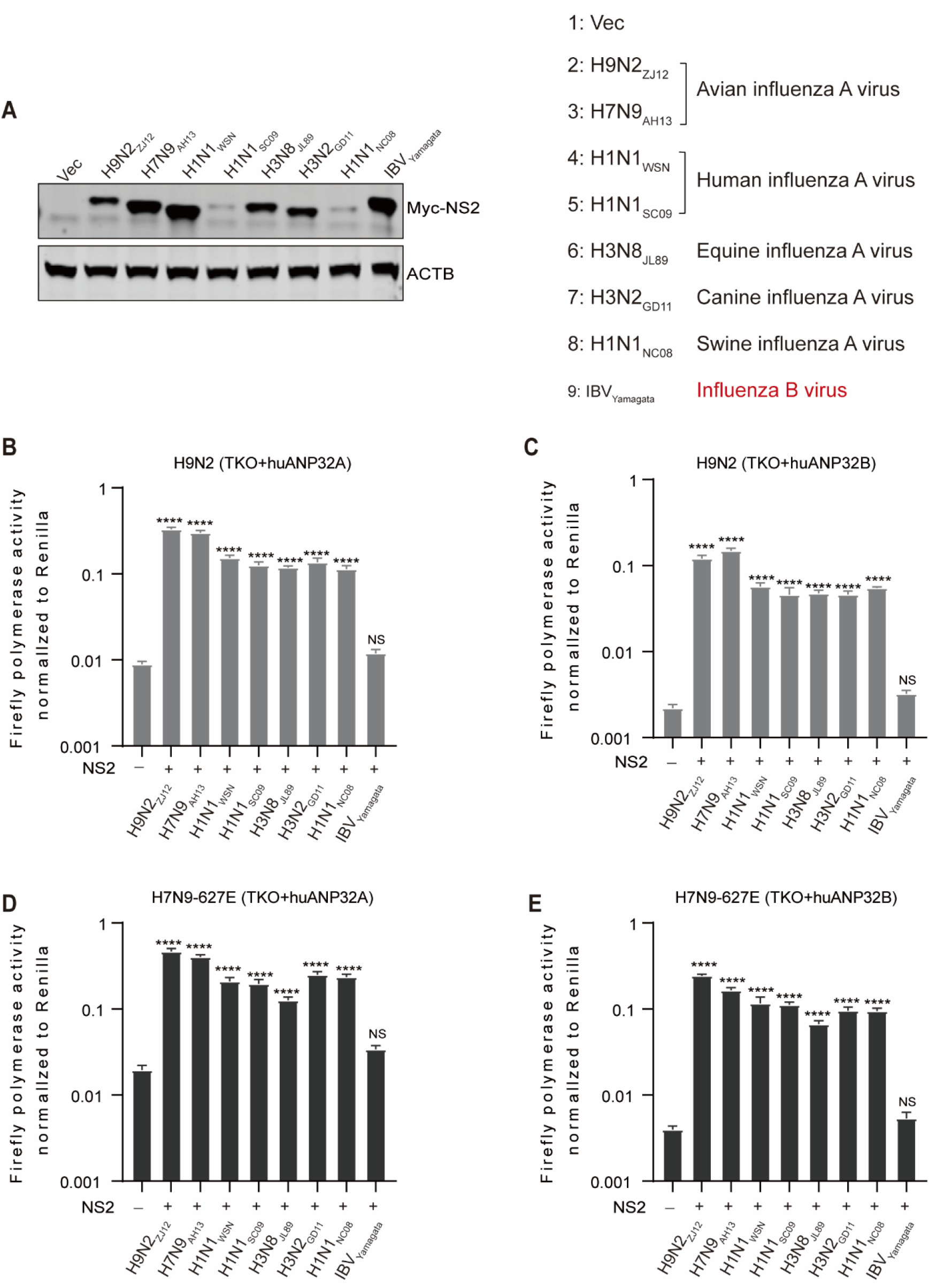
The avian-signature polymerase-enhancing function of NS2 is conserved (related to Fig.2) **A:** Expression levels of NS2 derived from influenza A viruses isolated from different host species and IBV. HEK293T cells were separately transfected with equal amounts of different Myc-tagged NS2 constructs. At 24 h post-transfection, cell lysates were prepared with RIPA lysis buffer and subjected to SDS-PAGE, followed by analysis with western blotting. **B:** Effect of different NS2 proteins on the H9N2 (B-C) and H7N9_AH13_ (PB2^627E^) (D-E) vPol activity when supported by huANP32A/B. Polymerase reconstitution assays were performed in TKO cells reconstituted with either huANP32A or huANP32B. Bars represent mean values of the replicates within one representative experiment (n = 3, ± SEM). Vec, empty vector control. Statistical significance was assessed using one-way ANOVA followed by Dunnett’s multiple comparisons test (\*\*\*\**p* < 0.0001; NS, non-significant).

**Figure S2.**
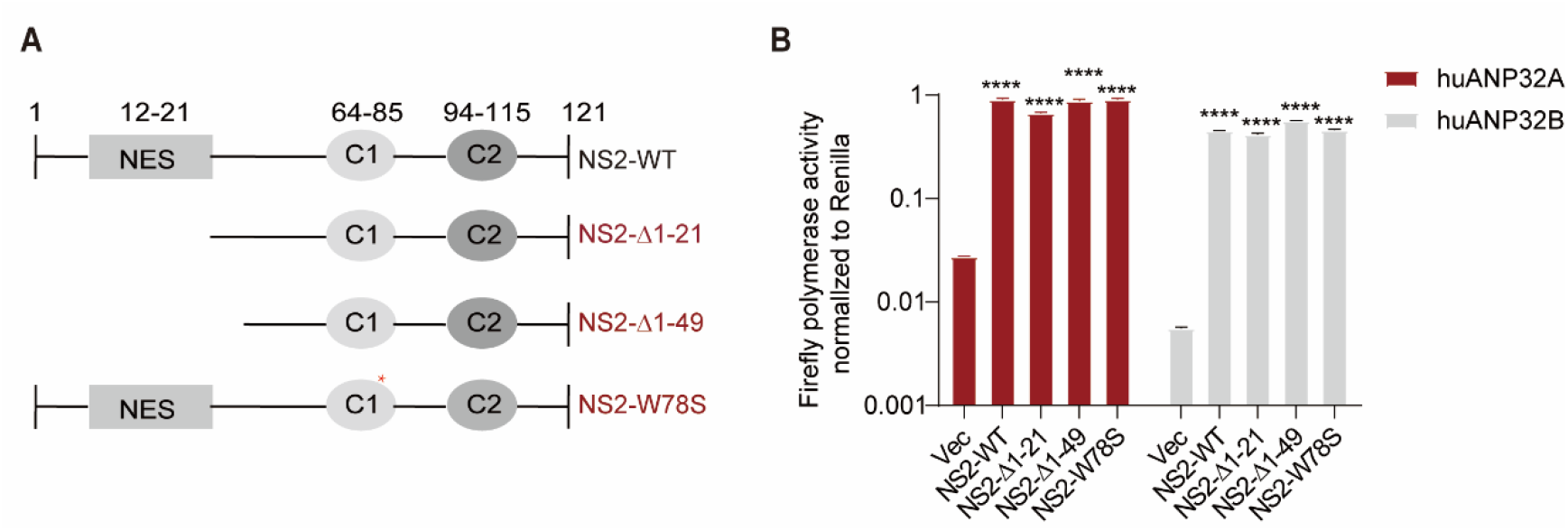
The function of NS2 in promoting the activity of avian-signature polymerase supported by mammalianANP32A/B is independent of its role in mediating nuclear export of vRNP (related to Fig.2) **A:** Diagram depicting mutants of NS2. **B:** Polymerase reconstitution assay in TKO cells to compare the effects of different NS2 mutants on H9N2 vPol activity when supported by either huANP32A or huANP32B. Bars represent mean values of the replicates within one representative experiment (n = 3, ± SD). Vec, empty vector control. Statistical significance was assessed using one-way ANOVA followed by Tukey’s multiple comparisons test (\*\*\*\**p* < 0.0001).

**Figure S3.**
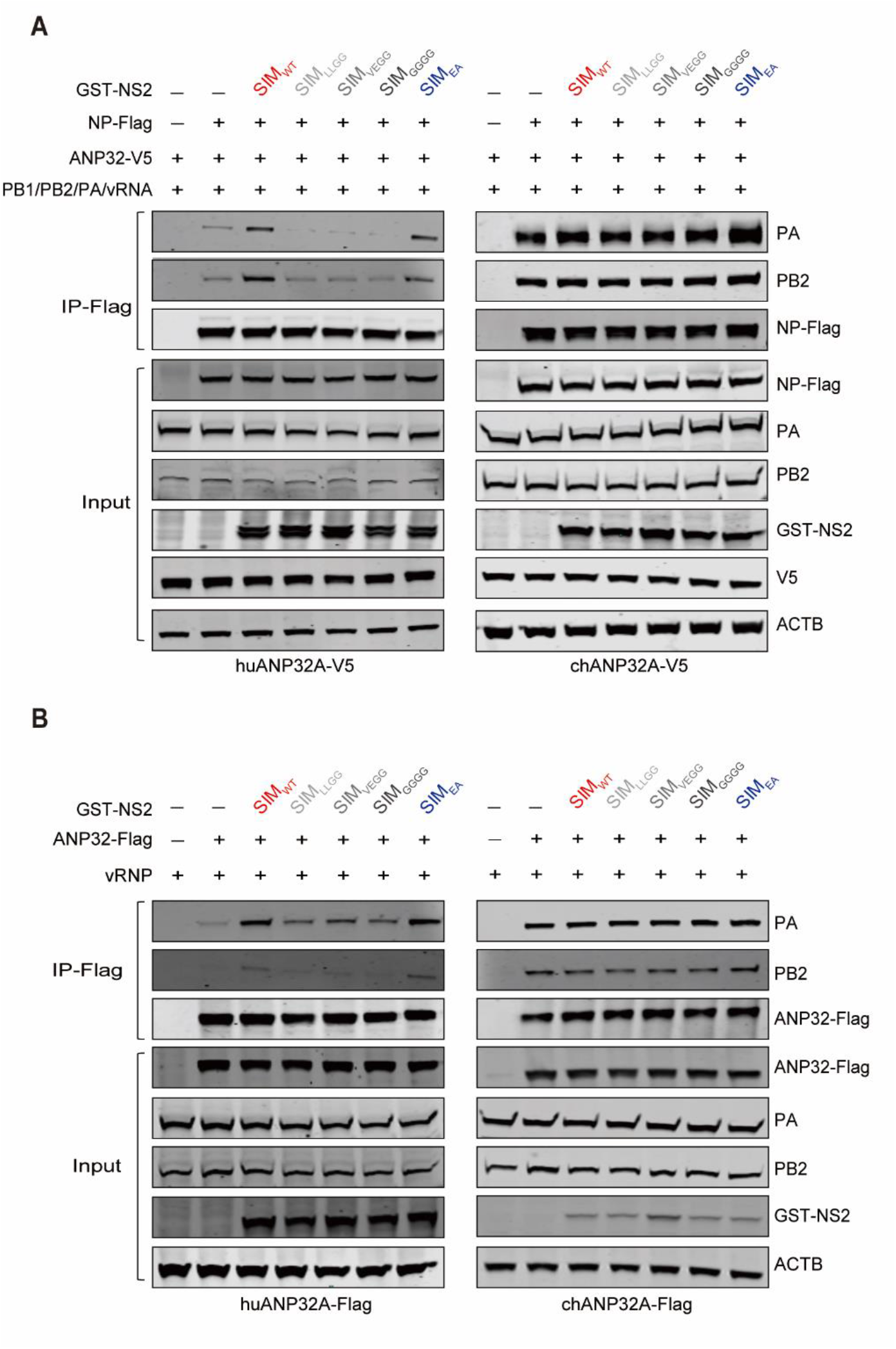
NS2 mutants in which the hydrophobic residues in the SIM have been replaced with glycines impair avian vRNP assembly as well as huANP32A-vRNP interactions (related to Fig.5) **A:** Effect of NS2 SIM mutations on H9N2 vRNP assembly in TKO cells reconstituted with either huANP32A or chANP32A. Experiments were performed as in Fig.4B, except that the 50 ng Myc-tagged NS2 was replaced with 3 μg GST-tagged NS2 constructs. **B:** Effect of NS2 SIM mutations on the H9N2 vRNP-ANP32 interactions in TKO cells reconstituted with either huANP32A or chANP32A. Experiments were performed as in Fig.4I, except that the 50 ng Myc-tagged NS2 was replaced with 3 μg GST-tagged NS2 constructs.

**Figure S4.**
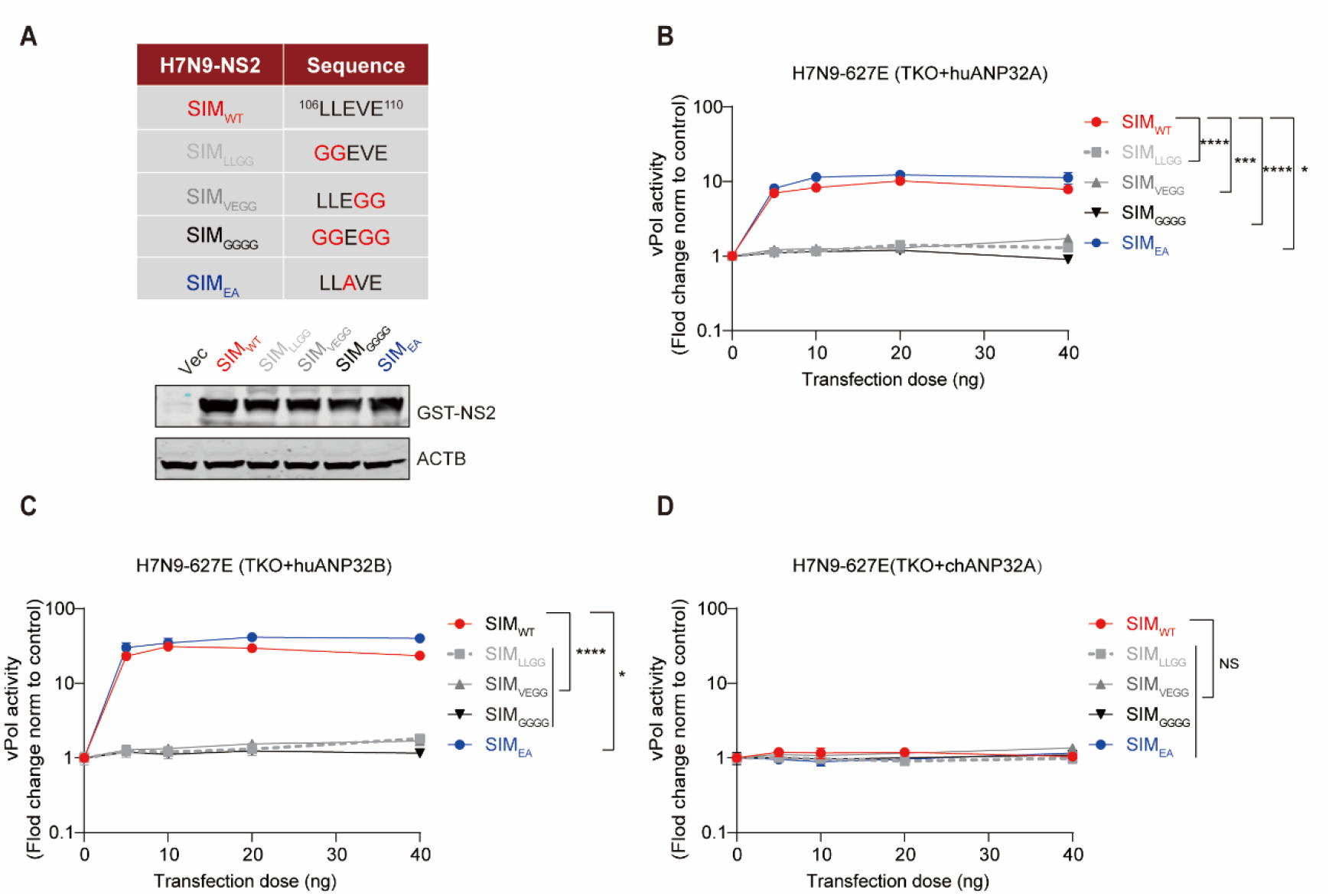
Impact of H7N9_AH13_ NS2 SIM sequence on the avian-signature vPol activity supported by huANP32A, huANP32B or chANP32A (related to Fig.5) **A:** Panel of constructed H7N9-NS2 mutants. The accompanying western blots show the impact of H7N9_AH13_ NS2 SIM changes on its protein expression levels. HEK293T cells were transfected with equal amounts of the indicted GST-tagged NS2 constructs. At 24 h post-transfection, cell lysates were prepared with RIPA lysis buffer and subjected to SDS-PAGE, followed by analysis by western blotting. **B:** Effect of different H7N9-NS2 mutants on the H7N9_AH13_ (PB2^627E^) vPol activity supported by huANP32A/B or chANP32A. vPol reconstitution assays were performed in TKO cells reconstituted separately with huANP32A(B), huANP32B(C) or chANP32A (D), together with increasing amounts of different NS2 constructs. For all assays, data were firefly activity normalized to *Renilla*, and plotted as fold change to empty vector (0 ng NS2 constructs). Bars represent mean values of the replicates within one representative experiment (n = 3, ± SD). Statistical significance was assessed using a two-way ANOVA (\**p* < 0.05; \*\*\**p* < 0.001; \*\*\*\**p* < 0.0001; NS, non-significant).

**Figure S5.**
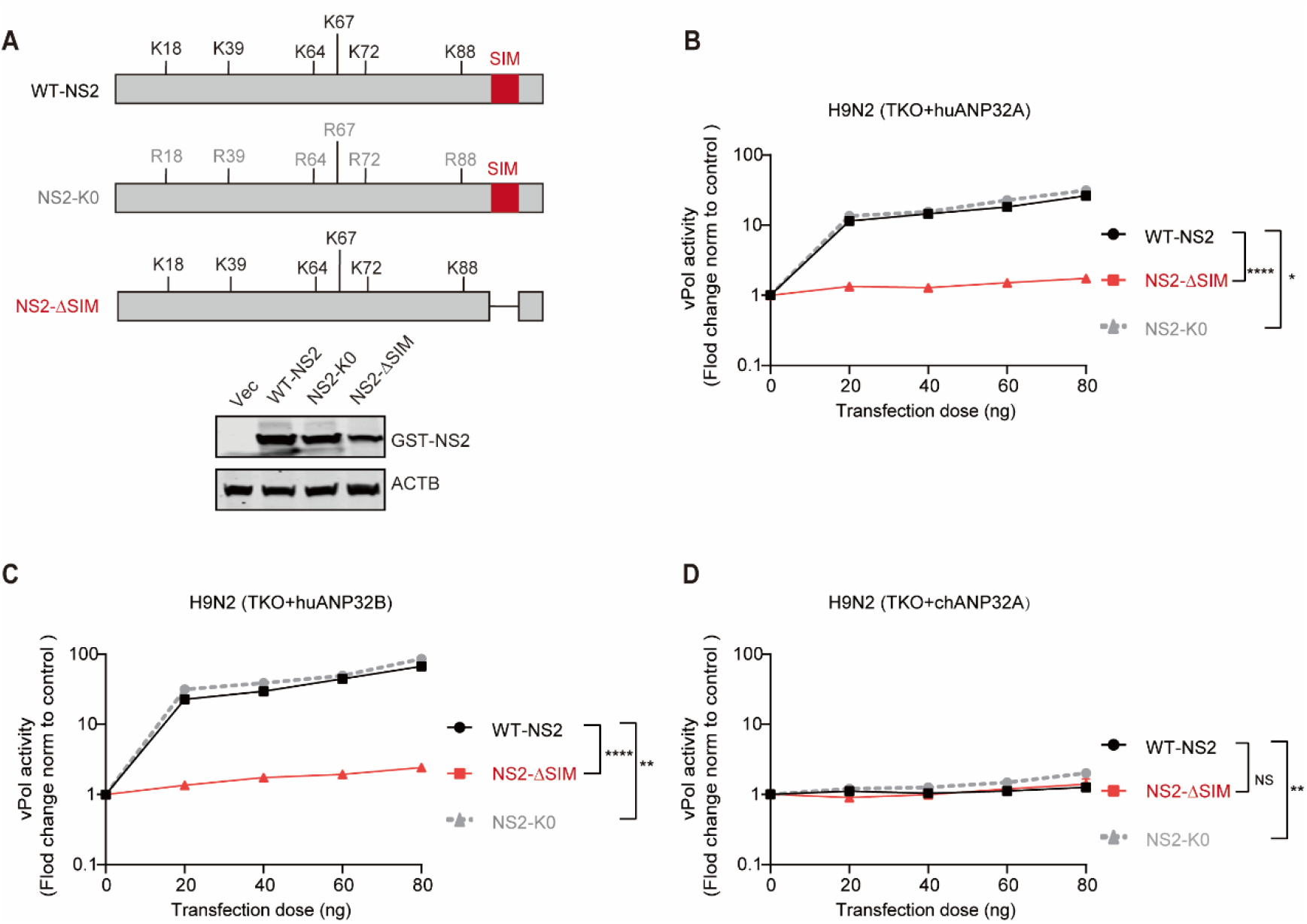
SUMOylation of NS2 is not required for its avian-signature polymerase-enhancing ability (related to Fig.5) **A:** Schematic representation of the NS2 mutants. The accompanying western blots show expression of GST-tagged NS2 constructs. HEK293T cells were transfected with equal amounts of indicted GST-tagged NS2 constructs. At 24 h post-transfection, cell lysates were prepared with RIPA lysis buffer and subjected to SDS-PAGE, followed by analysis by western blotting. **B-D:** Effect of different NS2 mutants on the H9N2 vPol activity supported by huANP32A/B or chANP32A. vPol reconstitution assays were performed in TKO cells reconstituted separately with huANP32A(B), huANP32B(C) or chANP32A (D), together with increasing amounts of different NS2 constructs. For all assays, data were firefly activity normalized to *Renilla*, and plotted as fold change to empty vector (0 ng NS2 constructs). Bars represent mean values of the replicates within one representative experiment (n = 3, ± SD). Statistical significance was assessed using a two-way ANOVA (\**p* < 0.05; \*\**p* < 0.01; \*\*\*\**p* < 0.0001; NS, non-significant).

**Figure S6.**
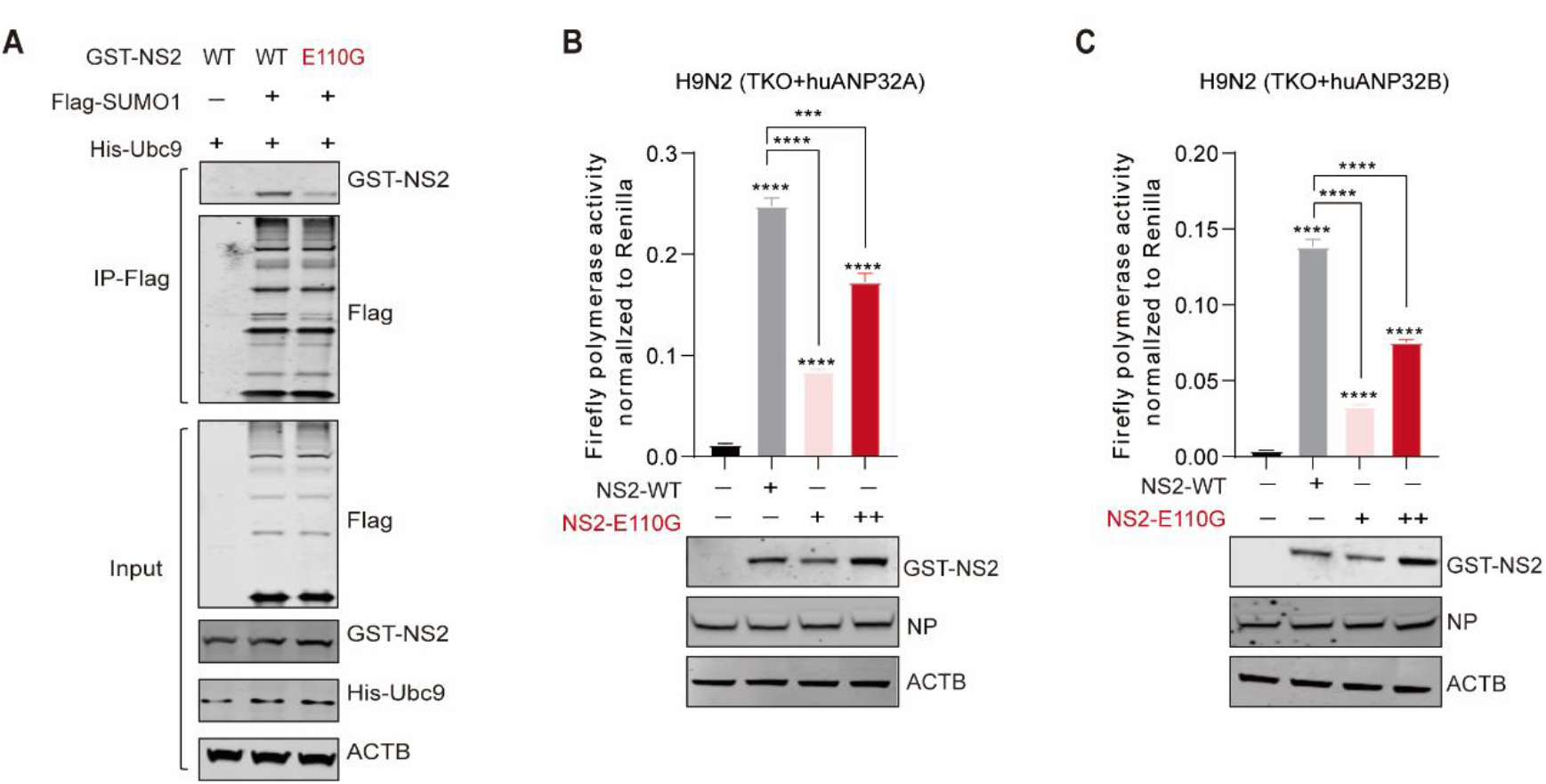
NS2 bearing a SIM E110G mutation defective in binding SUMO1-conjugated proteins impaired its ability to promote huANP32A/B-supported avian polymerase activity (related to Fig.6) **A:** SIM-E110G mutations resulted in the reduced binding of NS2 to SUMO1-conjugated proteins. HEK293T cells were transfected with GST-tagged NS2 constructs, together with co-expressed His-Ubc9, Flag-SUMO1 or empty vector control. 24 hours after transfection, lysed cells were subjected to immunoprecipitation with anti-Flag antibody, and coprecipitated GST-tagged NS2 constructs were detected using western blot analysis. **B-C:** SIM-E110G mutations impair the ability of NS2 to promote the H9N2 vPol activity supported by huANP32A (B) or huANP32B (C). Polymerase reconstitution assays were performed in TKO cells reconstituted with either huANP32A (B) or huANP32B (C), together with co-expressed empty vector, GST-tagged NS2-WT or NS2-E110G. The accompanying western blots show expression of GST-tagged NS2 constructs and vRNP component NP. In (B)–(C), bars represent mean values of the replicates within one representative experiment (n = 3, ± SD). Statistical significance was assessed using an unpaired Student’s *t-*tests (\**p* < 0.05; \*\**p* < 0.01; \*\*\**p* < 0.001; \*\*\*\**p* < 0.0001).

## RESOURCE AVAILABILITY

### Lead contact

Further information and requirements should be addressed to and will be fulfilled by lead contact Xiaojun Wang

### Materials availability

The materials used or generated in this study are available from the lead contact with a completed Materials Transfer Agreement.

### Data and code availability

- All data reported in this paper will be shared by the lead contact upon reasonable request.
- This paper does not report original code.
- Any additional information required to reanalyze the data presented in this work is available form the lead contact upon reasonable request.

## METHOD DETAILS

### Ethics statements

This study was conducted in strict accordance with the recommendations in the Guide for the Care and Use of Laboratory Animals of the Ministry of Science and Technology of the People’s Republic of China. The protocol for in vivo studies was approved by the Committee on the Ethics of Animal Experiments of the Harbin Veterinary Research Institute (HVRI) of the Chinese Academy of Agricultural Sciences (CAAS). 9-day-old embryonated chicken eggs and chicken blood were obtained from the National Poultry Laboratory Animal Resource Center.

### Biosafety statement and facility

All experiments using live H5N6 and H7N9 viruses were conducted within the enhanced animal biosafety level 3 (ABSL3+) facility at the HVRI of CAAS, which is approved for such use by the Ministry of Agriculture and Rural Affairs of China and the China National Accreditation Service for Conformity Assessment. Details of the facility and the biosafety and biosecurity measures used have been previously reported(Zhang et al., 2013b).

### Cells lines and constructs

HEK293T cells, TKO cells (*ANP32A, ANP32B* and *ANP32E* triple knockout HEK293T cell lines), MDCK cells, MDCK-chANP32A cells (MDCK cells stably expressing *chANP32A*) and primary chicken embryo fibroblast cells (CEF) were cultured in DMEM containing 10% FBS,1% penicillin-streptomycin (Pen-Strep) (Gibco). Preparation of TKO cells has been previously described(Zhang *et al*., 2020b). All cells were maintained at 37 °C, 5% CO _2_.

Preparation of the pCAGGS expression plasmids encoding the RNP proteins of H7N9 A/Anhui/01/2013(H7N9_AH13_), H9N2 avian influenza virus A/chicken/Zhejiang/B2013/2012 (H9N2_ZJ12_) and H1N1 human influenza virus A/WSN/1933 (WSN) has been described in previous studies(Zhang *et al*., 2020a; Zhang *et al*., 2019). pCAGGS expression plasmids encoding ANP32A and ANP32B from different species were kept in our laboratory and preparation was described in previous studies(Zhang *et al*., 2020a; Zhang *et al*., 2019; Zhang *et al*., 2020b). The ORFs of NS2 form influenza A virus isolated from different host species were cloned into the pCAGGS vector with a GST tag or Myc tag at the N-terminus. Site-directed mutants and deletion mutants of NS2 were generated with PCR and identified with DNA sequencing. The full-length ORFs of human SUMO1, SUMO2 and SUMO3 were cloned into the pCEF vector with a Flag tag at the N-terminus. To create the pCAGGS expression plasmids encoding for mutant SUMO1, SUMO2, and SUMO3, the required mutations were introduced with PCR using primers containing GG-to-AA mutations at the C-terminus of SUMO1/2/3. The ORFs of Ubc9 were cloned into the pCAGGS vectors with an N-terminal His_6_ tag. Development of the mini-genome reporters of influenza A and influenza B viruses and Renilla luciferase expression plasmids (pRL-TK) have been described previously(Zhang *et al*., 2020a; Zhang *et al*., 2019; Zhang *et al*., 2020b).

### vPol reconstitution assays

Cells plated in 24-well plates were cotransfected with pPolI-luc (40 ng), *Renilla* luciferase expression plasmid (5 ng), and four RNP expression plasmids pCAGGS-PB1 (20 ng), pCAGGS-PB2 (20 ng), pCAGGS-PA (10 ng), and pCAGGS–NP (40 ng), together with the indicated amount of the plasmids of interest. After 24 h of transfection, cells were lysed and luciferase activity was measured using the Dual-Glo Luciferase Assay system (Promega) with a Berthold Centro LB 960 Microplate Luminometer.

### Immunoprecipitations and GST pull-down

After transfection with the indicated plasmids for 24 h, transfected cells in a 25 cm^2^ flask were lysed with 1 mL lysis buffer (50 mM Hepes-NaOH [pH 7.9], 100 mM NaCl, 50 mM KCl, 0.25% NP-40, and 1 mM DTT). 100 μL cell lysate were saved as an input control and the rest of the cell lysate was immunoprecipitated with anti-Flag M2 Magnetic Beads (SIGMA-ALDRICH, M8823) at 4 °C overnight. The precipitated immune complexes were washed three times with lysis buffer and then eluted with 3 × Flag peptides. Immunoprecipitated proteins were then analyzed by western blotting.

For the GST pull-down assay, transfected cells in a 25 cm^2^ flask were lysed with 1 mL of lysis buffer as above. 100 μL of cell lysate was saved as an input control and the remaining cell lysate was incubated with Glutathione MagBeads (GenScript, L00327) overnight at 4°C. T he beads were then washed three times with 1 × PBS and boiled with sample loading buffer prior western blot analysis.

### Western blots

Western blot analysis was performed using the standard described methods^24^. The following antibodies were used: rabbit anti-GST (GenScript, A00097, 1:2000 for WB), rabbit anti-Flag (Sigma, F7425, 1:2000 for WB), rabbit anti-V5 (Proteintech, 14440-1-AP, 1:2000 for WB), mouse anti-His (Proteintech, 66005-1-Ig, 1:5000 for WB), rabbit anti-ACTB (Abclonal, AC026, 1:10000 for WB), mouse anti-ACTB (Abclonal, AC004, 1:2000 for WB), rabbit anti-Flag (Sigma-Aldrich, F7425, 1:5000 for WB), rabbit anti-influenza A virus PB2 (GeneTex, GTX125926, 1:5000 for WB), mouse anti-influenza A virus PA (prepared in our laboratory, 1:5000 for WB), mouse anti-influenza A virus PB2 (prepared in our laboratory, 1:1000 for WB), and mouse anti-influenza A virus NP (prepared in our laboratory, 1:1000 for WB).

### Influenza virus infection

Influenza A viruses containing NS2-SIM_WT_ or NS-SIM_E110G_ were made with the genome from H1N1 human influenza virus A/WSN/1933 (WSN, kindly provided by Dr Yoshihiro Kawaoka); H9N2 avian influenza virus A/chicken/Zhejiang/B2013/2012 (H9N2_ZJ12_, kindly provided by Dr Zejun Li); H7N9 avian influenza virus A/Chicken/Hunan/S1220/2017 (H7N9_HN17_) and H5N6 avian influenza virus A/duck/Guangdong/S1330/2016(H5N6_GD16_) were kept in our lab. Virus rescue was performed by transfecting TKO cells with a 12-plasmid rescue system. Briefly, TKO cells were co-transfected with eight pPol1 plasmids encoding all eight segments, as well as expression plasmids encoding RNP proteins and chANP32A. 24 h post-transfection, supernatants were harvested and injected into 9-day-old specific-pathogen-free embryonated eggs for virus propagation. Viral stocks were harvested after 48 h of incubation at 35 °C for and titrated in MDCK -chANP32A cell lines (MDCK cells stably expressing chANP32A) using standard methods. Viral genotypes were confirmed by DNA sequencing.

To determine virus growth curves, the indicated cells were infected with each virus diluted in opti-MEM and replaced 1-2 h later with opti-MEM supplemented with 0-2μg/ml TPCK-treated trypsin. Cell supernatants were harvested at the indicated time points post-infection and the viral titers were determined in MDCK-chANP32A cells using TCID_50_ assay.

### Mouse study

To determine the viral replication in mice, the indicated dose of H5N6-SIM_WT_ or H5N6-SIM_E110G_ was diluted in PBS and used for intranasal inoculation of five female BALB/c mice. The mice were euthanized for the collection of nasal turbinate and lungs on day 3 p.i, day 5 p.i and day 7 p.i, respectively. Viral titers were then determined in MDCK-chANP32A cells using TCID_50_ assays.

To determine the MLD_50_ value, groups of five BALB/c mice were intranasally inoculated with 10^1^-10^6^ EID_50_ of virus in a volume of 50 μL. Then the inoculated mice were monitored for mortality for two weeks and the MLD_50_ values were calculated by the Reed and Muench method s(Reed, 1938).

### Chicken study

To determine the virulence of wild type H5N6 and their mutants in chickens, an Intravenous Pathogenicity Index (IVPI) was performed according to the method described in the “Diagnostic Manual for Avian Influenza”. Ten 6-week-old chickens were inoculated intravenously with the virus, after which the mortality was monitored for 10 days. To determine the ability of H5N6 virus to replicate in chickens, three chickens were inoculated i.n. with 10^6^ EID_50_ of in a 0.1 ml volume. The chickens were killed on day 3 p.i., and their organs, including brains, lungs, kidneys, spleens, pancreas, duodenums, liver, tracheae, and ceca were collected for viral titration in MDCK-chANP32A cells using TCID_50_ assays.

### Statistical analysis

Quantitative data are presented as means ± SD as indicated in the figure legends. Statistical differences were analyzed with an unpaired Student’s *t*-test, two-way-ANOVA or one-way ANOVA followed by a Dunnett’s multiple comparisons test or a Tukey’s multiple comparisons test using GraphPad Prism 7.0 software. Statistical parameters are reported in the figures and figure legends (NS, not significant; *p* > 0.05; *, *p* < 0.05; **, *p* < 0.01; ***, *p* < 0.001; ****, *p* < 0.0001).

## REFERENCE

Akarsu, H., Burmeister, W.P., Petosa, C., Petit, I., Muller, C.W., Ruigrok, R.W., and Baudin, F. (2003). Crystal structure of the M1 protein-binding domain of the influenza A virus nuclear export protein (NEP/NS2). EMBO J 22, 4646–4655. 10.1093/emboj/cdg449.

Almond, J.W. (1977). A single gene determines the host range of influenza virus. Nature 270, 617–618. 10.1038/270617a0.

Baker, S.F., Ledwith, M.P., and Mehle, A. (2018). Differential Splicing of ANP32A in Birds Alters Its Ability to Stimulate RNA Synthesis by Restricted Influenza Polymerase. Cell Rep 24, 2581–2588 e2584. 10.1016/j.celrep.2018.08.012.

Belser, J.A., Davis, C.T., Balish, A., Edwards, L.E., Zeng, H., Maines, T.R., Gustin, K.M., Martinez, I.L., Fasce, R., Cox, N.J., et al. (2013). Pathogenesis, transmissibility, and ocular tropism of a highly pathogenic avian influenza A (H7N3) virus associated with human conjunctivitis. J Virol 87, 5746–5754. 10.1128/JVI.00154-13.

Bi, Y., Chen, Q., Wang, Q., Chen, J., Jin, T., Wong, G., Quan, C., Liu, J., Wu, J., Yin, R., et al. (2016). Genesis, Evolution and Prevalence of H5N6 Avian Influenza Viruses in China. Cell Host Microbe 20, 810–821. 10.1016/j.chom.2016.10.022.

Bi, Z., Ye, H., Wang, X., Fang, A., Yu, T., Yan, L., and Zhou, J. (2019). Insights into species-specific regulation of ANP32A on the mammalian-restricted influenza virus polymerase activity. Emerg Microbes Infect 8, 1465–1478. 10.1080/22221751.2019.1676625.

Chen, H., Yuan, H., Gao, R., Zhang, J., Wang, D., Xiong, Y., Fan, G., Yang, F., Li, X., Zhou, J., et al. (2014). Clinical and epidemiological characteristics of a fatal case of avian influenza A H10N8 virus infection: a descriptive study. Lancet 383, 714–721. 10.1016/S0140-6736(14)60111-2.

Cui, P., Zeng, X., Li, X., Li, Y., Shi, J., Zhao, C., Qu, Z., Wang, Y., Guo, J., Gu, W., et al. (2022). Genetic and biological characteristics of the globally circulating H5N8 avian influenza viruses and the protective efficacy offered by the poultry vaccine currently used in China. Sci China Life Sci 65, 795–808. 10.1007/s11427-021-2025-y.

Domingues, P., Eletto, D., Magnus, C., Turkington, H.L., Schmutz, S., Zagordi, O., Lenk, M., Beer, M., Stertz, S., and Hale, B.G. (2019). Profiling host ANP32A splicing landscapes to predict influenza A virus polymerase adaptation. Nat Commun 10, 3396. 10.1038/s41467-019-11388-2.

Domingues, P., Golebiowski, F., Tatham, M.H., Lopes, A.M., Taggart, A., Hay, R.T., and Hale, B.G. (2015). Global Reprogramming of Host SUMOylation during Influenza Virus Infection. Cell Rep 13, 1467–1480. 10.1016/j.celrep.2015.10.001.

Domingues, P., and Hale, B.G. (2017). Functional Insights into ANP32A-Dependent Influenza A Virus Polymerase Host Restriction. Cell Rep 20, 2538–2546. 10.1016/j.celrep.2017.08.061.

Eisfeld, A.J., Neumann, G., and Kawaoka, Y. (2015). At the centre: influenza A virus ribonucleoproteins. Nat Rev Microbiol 13, 28–41. 10.1038/nrmicro3367.

Fodor, E., and Te Velthuis, A.J.W. (2019). Structure and Function of the Influenza Virus Transcription and Replication Machinery. Cold Spring Harb Perspect Med. 10.1101/cshperspect.a038398.

Gabriel, G., Dauber, B., Wolff, T., Planz, O., Klenk, H.D., and Stech, J. (2005). The viral polymerase mediates adaptation of an avian influenza virus to a mammalian host. Proc Natl Acad Sci U S A 102, 18590–18595. 10.1073/pnas.0507415102.

Gao, Y., Zhang, Y., Shinya, K., Deng, G., Jiang, Y., Li, Z., Guan, Y., Tian, G., Li, Y., Shi, J., et al. (2009). Identification of amino acids in HA and PB2 critical for the transmission of H5N1 avian influenza viruses in a mammalian host. PLoS Pathog 5, e1000709. 10.1371/journal.ppat.1000709.

Geiss-Friedlander, R., and Melchior, F. (2007). Concepts in sumoylation: a decade on. Nat Rev Mol Cell Biol 8, 947–956. 10.1038/nrm2293.

Gu, W., Shi, J., Cui, P., Yan, C., Zhang, Y., Wang, C., Zhang, Y., Xing, X., Zeng, X., Liu, L., et al. (2022). Novel H5N6 reassortants bearing the clade 2.3.4.4b HA gene of H5N8 virus have been detected in poultry and caused multiple human infections in China. Emerg Microbes Infect 11, 1174–1185. 10.1080/22221751.2022.2063076.

Guan, L., Shi, J., Kong, X., Ma, S., Zhang, Y., Yin, X., He, X., Liu, L., Suzuki, Y., Li, C., et al. (2019). H3N2 avian influenza viruses detected in live poultry markets in China bind to human-type receptors and transmit in guinea pigs and ferrets. Emerg Microbes Infect 8, 1280–1290. 10.1080/22221751.2019.1660590.

Guo, J., Chen, J., Li, Y., Li, Y., Deng, G., Shi, J., Liu, L., Chen, H., and Li, X. (2022). SUMOylation of Matrix Protein M1 and Filamentous Morphology Collectively Contribute to the Replication and Virulence of Highly Pathogenic H5N1 Avian Influenza Viruses in Mammals. J Virol 96, e0163021. 10.1128/JVI.01630-21.

Guo, J., Wang, Y., Zhao, C., Gao, X., Zhang, Y., Li, J., Wang, M., Zhang, H., Liu, W., Wang, C., et al. (2021). Molecular characterization, receptor binding property, and replication in chickens and mice of H9N2 avian influenza viruses isolated from chickens, peafowls, and wild birds in eastern China. Emerg Microbes Infect 10, 2098–2112. 10.1080/22221751.2021.1999778.

Han, Q., Chang, C., Li, L., Klenk, C., Cheng, J., Chen, Y., Xia, N., Shu, Y., Chen, Z., Gabriel, G., et al. (2014). Sumoylation of influenza A virus nucleoprotein is essential for intracellular trafficking and virus growth. J Virol 88, 9379–9390. 10.1128/JVI.00509-14.

Hatta, M., Gao, P., Halfmann, P., and Kawaoka, Y. (2001). Molecular basis for high virulence of Hong Kong H5N1 influenza A viruses. Science 293, 1840–1842. 10.1126/science.1062882.

Hecker, C.M., Rabiller, M., Haglund, K., Bayer, P., and Dikic, I. (2006). Specification of SUMO1- and SUMO2-interacting motifs. J Biol Chem 281, 16117–16127. 10.1074/jbc.M512757200.

Jonges, M., Welkers, M.R., Jeeninga, R.E., Meijer, A., Schneeberger, P., Fouchier, R.A., de Jong, M.D., and Koopmans, M. (2014). Emergence of the virulence-associated PB2 E627K substitution in a fatal human case of highly pathogenic avian influenza virus A(H7N7) infection as determined by Illumina ultra-deep sequencing. J Virol 88, 1694–1702. 10.1128/JVI.02044-13.

Labadie, K., Dos Santos Afonso, E., Rameix-Welti, M.A., van der Werf, S., and Naffakh, N. (2007). Host-range determinants on the PB2 protein of influenza A viruses control the interaction between the viral polymerase and nucleoprotein in human cells. Virology 362, 271–282. 10.1016/j.virol.2006.12.027.

Le, Q.M., Ito, M., Muramoto, Y., Hoang, P.V., Vuong, C.D., Sakai-Tagawa, Y., Kiso, M., Ozawa, M., Takano, R., and Kawaoka, Y. (2010). Pathogenicity of highly pathogenic avian H5N1 influenza A viruses isolated from humans between 2003 and 2008 in northern Vietnam. J Gen Virol 91, 2485–2490. 10.1099/vir.0.021659-0.

Li, C., and Chen, H. (2021). H7N9 Influenza Virus in China. Cold Spring Harb Perspect Med 11. 10.1101/cshperspect.a038349.

Li, J., Liang, L., Jiang, L., Wang, Q., Wen, X., Zhao, Y., Cui, P., Zhang, Y., Wang, G., Li, Q., et al. (2021). Viral RNA-binding ability conferred by SUMOylation at PB1 K612 of influenza A virus is essential for viral pathogenesis and transmission. PLoS Pathog 17, e1009336. 10.1371/journal.ppat.1009336.

Li, Q., Zhou, L., Zhou, M., Chen, Z., Li, F., Wu, H., Xiang, N., Chen, E., Tang, F., Wang, D., et al. (2014a). Epidemiology of human infections with avian influenza A(H7N9) virus in China. N Engl J Med 370, 520–532. 10.1056/NEJMoa1304617.

Li, X., Shi, J., Guo, J., Deng, G., Zhang, Q., Wang, J., He, X., Wang, K., Chen, J., Li, Y., et al. (2014b). Genetics, receptor binding property, and transmissibility in mammals of naturally isolated H9N2 Avian Influenza viruses. PLoS Pathog 10, e1004508. 10.1371/journal.ppat.1004508.

Li, Y., Shi, J., Zhong, G., Deng, G., Tian, G., Ge, J., Zeng, X., Song, J., Zhao, D., Liu, L., et al. (2010). Continued evolution of H5N1 influenza viruses in wild birds, domestic poultry, and humans in China from 2004 to 2009. J Virol 84, 8389–8397. 10.1128/JVI.00413-10.

Liang, L., Jiang, L., Li, J., Zhao, Q., Wang, J., He, X., Huang, S., Wang, Q., Zhao, Y., Wang, G., et al. (2019). Low Polymerase Activity Attributed to PA Drives the Acquisition of the PB2 E627K Mutation of H7N9 Avian Influenza Virus in Mammals. MBio 10. 10.1128/mBio.01162-19.

Lin, D.Y., Huang, Y.S., Jeng, J.C., Kuo, H.Y., Chang, C.C., Chao, T.T., Ho, C.C., Chen, Y.C., Lin, T.P., Fang, H.I., et al. (2006). Role of SUMO-interacting motif in Daxx SUMO modification, subnuclear localization, and repression of sumoylated transcription factors. Mol Cell 24, 341–354. 10.1016/j.molcel.2006.10.019.

Long, J.S., Giotis, E.S., Moncorge, O., Frise, R., Mistry, B., James, J., Morisson, M., Iqbal, M., Vignal, A., Skinner, M.A., and Barclay, W.S. (2016). Species difference in ANP32A underlies influenza A virus polymerase host restriction. Nature 529, 101–104. 10.1038/nature16474.

Long, J.S., Idoko-Akoh, A., Mistry, B., Goldhill, D., Staller, E., Schreyer, J., Ross, C., Goodbourn, S., Shelton, H., Skinner, M.A., et al. (2019a). Species specific differences in use of ANP32 proteins by influenza A virus. Elife 8. 10.7554/eLife.45066.

Long, J.S., Mistry, B., Haslam, S.M., and Barclay, W.S. (2019b). Host and viral determinants of influenza A virus species specificity. Nat Rev Microbiol 17, 67–81. 10.1038/s41579-018-0115-z.

Manz, B., Brunotte, L., Reuther, P., and Schwemmle, M. (2012). Adaptive mutations in NEP compensate for defective H5N1 RNA replication in cultured human cells. Nat Commun 3, 802. 10.1038/ncomms1804.

Manz, B., de Graaf, M., Mogling, R., Richard, M., Bestebroer, T.M., Rimmelzwaan, G.F., and Fouchier, R.A.M. (2016). Multiple Natural Substitutions in Avian Influenza A Virus PB2 Facilitate Efficient Replication in Human Cells. J Virol 90, 5928–5938. 10.1128/JVI.00130-16.

Martinat, C., Cormier, A., Tobaly-Tapiero, J., Palmic, N., Casartelli, N., Mahboubi, B., Coggins, S.A., Buchrieser, J., Persaud, M., Diaz-Griffero, F., et al. (2021). SUMOylation of SAMHD1 at Lysine 595 is required for HIV-1 restriction in non-cycling cells. Nat Commun 12, 4582. 10.1038/s41467-021-24802-5.

Mehle, A., and Doudna, J.A. (2008). An inhibitory activity in human cells restricts the function of an avian-like influenza virus polymerase. Cell Host Microbe 4, 111–122. 10.1016/j.chom.2008.06.007.

Mok, C.K., Lee, H.H., Lestra, M., Nicholls, J.M., Chan, M.C., Sia, S.F., Zhu, H., Poon, L.L., Guan, Y., and Peiris, J.S. (2014). Amino acid substitutions in polymerase basic protein 2 gene contribute to the pathogenicity of the novel A/H7N9 influenza virus in mammalian hosts. J Virol 88, 3568–3576. 10.1128/JVI.02740-13.

Neumann, G., Chen, H., Gao, G.F., Shu, Y., and Kawaoka, Y. (2010). H5N1 influenza viruses: outbreaks and biological properties. Cell Res 20, 51–61. 10.1038/cr.2009.124.

Neumann, G., Hughes, M.T., and Kawaoka, Y. (2000). Influenza A virus NS2 protein mediates vRNP nuclear export through NES-independent interaction with hCRM1. EMBO J 19, 6751–6758. 10.1093/emboj/19.24.6751.

O’Neill, R.E., Talon, J., and Palese, P. (1998). The influenza virus NEP (NS2 protein) mediates the nuclear export of viral ribonucleoproteins. EMBO J 17, 288–296. 10.1093/emboj/17.1.288.

Office International des Epizooties, P., 2011 Manual of diagnostic tests and vaccines for terrestrial animal.

Pal, S., Santos, A., Rosas, J.M., Ortiz-Guzman, J., and Rosas-Acosta, G. (2011). Influenza A virus interacts extensively with the cellular SUMOylation system during infection. Virus Res 158, 12–27. 10.1016/j.virusres.2011.02.017.

Pan, Y., Cui, S., Sun, Y., Zhang, X., Ma, C., Shi, W., Peng, X., Lu, G., Zhang, D., Liu, Y., et al. (2018). Human infection with H9N2 avian influenza in northern China. Clin Microbiol Infect 24, 321–323. 10.1016/j.cmi.2017.10.026.

Paterson, D., and Fodor, E. (2012). Emerging roles for the influenza A virus nuclear export protein (NEP). PLoS Pathog 8, e1003019. 10.1371/journal.ppat.1003019.

Peacock, T.P., Swann, O.C., Salvesen, H.A., Staller, E., Leung, P.B., Goldhill, D.H., Zhou, H., Lillico, S.G., Whitelaw, C.B.A., Long, J.S., and Barclay, W.S. (2020). Swine ANP32A Supports Avian Influenza Virus Polymerase. J Virol 94. 10.1128/JVI.00132-20.

Perez, J.T., Varble, A., Sachidanandam, R., Zlatev, I., Manoharan, M., Garcia-Sastre, A., and tenOever, B.R. (2010). Influenza A virus-generated small RNAs regulate the switch from transcription to replication. Proc Natl Acad Sci U S A 107, 11525–11530. 10.1073/pnas.1001984107.

Reed, L.J. (1938). A simple method of estimating fifty percent endpoints. Am J Hyg 27.

Resa-Infante, P., Jorba, N., Coloma, R., and Ortin, J. (2011). The influenza virus RNA synthesis machine: advances in its structure and function. RNA Biol 8, 207–215. 10.4161/rna.8.2.14513.

Robb, N.C., Smith, M., Vreede, F.T., and Fodor, E. (2009). NS2/NEP protein regulates transcription and replication of the influenza virus RNA genome. J Gen Virol 90, 1398–1407. 10.1099/vir.0.009639-0.

Rodriguez, M.S., Dargemont, C., and Hay, R.T. (2001). SUMO-1 conjugation in vivo requires both a consensus modification motif and nuclear targeting. J Biol Chem 276, 12654–12659. 10.1074/jbc.M009476200.

Sampson, D.A., Wang, M., and Matunis, M.J. (2001). The small ubiquitin-like modifier-1 (SUMO-1) consensus sequence mediates Ubc9 binding and is essential for SUMO-1 modification. J Biol Chem 276, 21664–21669. 10.1074/jbc.M100006200.

Shi, J., Deng, G., Kong, H., Gu, C., Ma, S., Yin, X., Zeng, X., Cui, P., Chen, Y., Yang, H., et al. (2017). H7N9 virulent mutants detected in chickens in China pose an increased threat to humans. Cell Res 27, 1409–1421. 10.1038/cr.2017.129.

Shi, J., Deng, G., Ma, S., Zeng, X., Yin, X., Li, M., Zhang, B., Cui, P., Chen, Y., Yang, H., et al. (2018). Rapid Evolution of H7N9 Highly Pathogenic Viruses that Emerged in China in 2017. Cell Host Microbe 24, 558–568 e557. 10.1016/j.chom.2018.08.006.

Song, J., Durrin, L.K., Wilkinson, T.A., Krontiris, T.G., and Chen, Y. (2004). Identification of a SUMO-binding motif that recognizes SUMO-modified proteins. Proc Natl Acad Sci U S A 101, 14373–14378. 10.1073/pnas.0403498101.

Song, W., and Qin, K. (2020). Human-infecting influenza A (H9N2) virus: A forgotten potential pandemic strain? Zoonoses Public Health 67, 203–212. 10.1111/zph.12685.

Staller, E., Sheppard, C.M., Neasham, P.J., Mistry, B., Peacock, T.P., Goldhill, D.H., Long, J.S., and Barclay, W.S. (2019). ANP32 proteins are essential for influenza virus replication in human cells. J Virol. 10.1128/JVI.00217-19.

Steel, J., Lowen, A.C., Mubareka, S., and Palese, P. (2009). Transmission of influenza virus in a mammalian host is increased by PB2 amino acids 627K or 627E/701N. PLoS Pathog 5, e1000252. 10.1371/journal.ppat.1000252.

Su, S., Gu, M., Liu, D., Cui, J., Gao, G.F., Zhou, J., and Liu, X. (2017). Epidemiology, Evolution, and Pathogenesis of H7N9 Influenza Viruses in Five Epidemic Waves since 2013 in China. Trends Microbiol 25, 713–728. 10.1016/j.tim.2017.06.008.

Subbarao, E.K., London, W., and Murphy, B.R. (1993). A single amino acid in the PB2 gene of influenza A virus is a determinant of host range. J Virol 67, 1761–1764. 10.1128/JVI.67.4.1761-1764.1993.

Sugiyama, K., Kawaguchi, A., Okuwaki, M., and Nagata, K. (2015). pp32 and APRIL are host cell-derived regulators of influenza virus RNA synthesis from cRNA. Elife 4. 10.7554/eLife.08939.

Van Hoeven, N., Pappas, C., Belser, J.A., Maines, T.R., Zeng, H., Garcia-Sastre, A., Sasisekharan, R., Katz, J.M., and Tumpey, T.M. (2009). Human HA and polymerase subunit PB2 proteins confer transmission of an avian influenza virus through the air. Proc Natl Acad Sci U S A 106, 3366–3371. 10.1073/pnas.0813172106.

Wandzik, J.M., Kouba, T., and Cusack, S. (2021). Structure and Function of Influenza Polymerase. Cold Spring Harb Perspect Med 11. 10.1101/cshperspect.a038372.

Wang, G., Zhao, Y., Zhou, Y., Jiang, L., Liang, L., Kong, F., Yan, Y., Wang, X., Wang, Y., Wen, X., et al. (2022). PIAS1-mediated SUMOylation of influenza A virus PB2 restricts viral replication and virulence. PLoS Pathog 18, e1010446. 10.1371/journal.ppat.1010446.

Wei, S.H., Yang, J.R., Wu, H.S., Chang, M.C., Lin, J.S., Lin, C.Y., Liu, Y.L., Lo, Y.C., Yang, C.H., Chuang, J.H., et al. (2013). Human infection with avian influenza A H6N1 virus: an epidemiological analysis. Lancet Respir Med 1, 771–778. 10.1016/S2213-2600(13)70221-2.

Wu, C.Y., Jeng, K.S., and Lai, M.M. (2011). The SUMOylation of matrix protein M1 modulates the assembly and morphogenesis of influenza A virus. J Virol 85, 6618–6628. 10.1128/JVI.02401-10.

Xu, K., Klenk, C., Liu, B., Keiner, B., Cheng, J., Zheng, B.J., Li, L., Han, Q., Wang, C., Li, T., et al. (2011). Modification of nonstructural protein 1 of influenza A virus by SUMO1. J Virol 85, 1086–1098. 10.1128/JVI.00877-10.

Yin, X., Deng, G., Zeng, X., Cui, P., Hou, Y., Liu, Y., Fang, J., Pan, S., Wang, D., Chen, X., et al. (2021). Genetic and biological properties of H7N9 avian influenza viruses detected after application of the H7N9 poultry vaccine in China. PLoS Pathog 17, e1009561. 10.1371/journal.ppat.1009561.

Zhang, H., Li, H., Wang, W., Wang, Y., Han, G.Z., Chen, H., and Wang, X. (2020a). A unique feature of swine ANP32A provides susceptibility to avian influenza virus infection in pigs. PLoS Pathog 16, e1008330. 10.1371/journal.ppat.1008330.

Zhang, H., Zhang, Z., Wang, Y., Wang, M., Wang, X., Zhang, X., Ji, S., Du, C., Chen, H., and Wang, X. (2019). Fundamental Contribution and Host Range Determination of ANP32A and ANP32B in Influenza A Virus Polymerase Activity. J Virol 93. 10.1128/JVI.00174-19.

Zhang, Q., Shi, J., Deng, G., Guo, J., Zeng, X., He, X., Kong, H., Gu, C., Li, X., Liu, J., et al. (2013a). H7N9 influenza viruses are transmissible in ferrets by respiratory droplet. Science 341, 410–414. 10.1126/science.1240532.

Zhang, Y., Zhang, Q., Kong, H., Jiang, Y., Gao, Y., Deng, G., Shi, J., Tian, G., Liu, L., Liu, J., et al. (2013b). H5N1 hybrid viruses bearing 2009/H1N1 virus genes transmit in guinea pigs by respiratory droplet. Science 340, 1459–1463. 10.1126/science.1229455.

Zhang, Z., Zhang, H., Xu, L., Guo, X., Wang, W., Ji, Y., Lin, C., Wang, Y., and Wang, X. (2020b). Selective usage of ANP32 proteins by influenza B virus polymerase: Implications in determination of host range. PLoS Pathog 16, e1008989. 10.1371/journal.ppat.1008989.

Zhao, D., Liang, L., Li, Y., Jiang, Y., Liu, L., and Chen, H. (2012). Phylogenetic and pathogenic analyses of avian influenza A H5N1 viruses isolated from poultry in Vietnam. PLoS One 7, e50959. 10.1371/journal.pone.0050959.

Zhao, Q., Xie, Y., Zheng, Y., Jiang, S., Liu, W., Mu, W., Liu, Z., Zhao, Y., Xue, Y., and Ren, J. (2014). GPS-SUMO: a tool for the prediction of sumoylation sites and SUMO-interaction motifs. Nucleic Acids Res 42, W325–330. 10.1093/nar/gku383.

Zhou, J., Wang, D., Gao, R., Zhao, B., Song, J., Qi, X., Zhang, Y., Shi, Y., Yang, L., Zhu, W., et al. (2013). Biological features of novel avian influenza A (H7N9) virus. Nature 499, 500–503. 10.1038/nature12379.

